# Transfer seeding reveals divergent activation ensembles across tyrosine kinases

**DOI:** 10.64898/2026.06.26.734786

**Authors:** Yanchen Zhu, Matteo T. Degiacomi, Antonia S. J. S. Mey, Sukrit Singh

**Author notes:** Correspondence (M.T.D); (A.S.J.S.M.); (S.S.).

## Abstract

Protein kinase activation involves coordinated rearrangements of conserved regulatory elements, including the Asp–Phe–Gly (DFG) motif and activation loop (A-loop), but whether tyrosine kinases follow a universal activation mechanism remains unclear. This motivates the need for a common protocol to generate comparable conformational ensembles despite uneven experimental structural coverage across kinases. Here we combine over 8.3 milliseconds of distributed unbiased molecular dynamics simulations with Markov state models (MSMs) to compare the conformational landscapes of the ABL, EGFR and MET kinase domains. To expand conformational coverage while retaining unbiased dynamics, we use a transfer seeding strategy that steers AlphaFold2 models derived from homologous templates to sample MET conformational states absent from available experimental databases. Our MSMs resolve metastable conformations not represented in the experimental structural set, including low-population DFG-inter intermediates along the MET activation pathway. We find that activation pathways are kinase-specific rather than universal. A-loop rearrangements contribute to the rate-limiting steps in ABL and MET, whereas EGFR is rate limited by a local rearrangement of the DFG motif. Building on established DFG geometry nomenclature, ensemble level characterization of the A-loop further distinguishes metastable conformations that share the same DFG geometry and reveal kinase-specific differences in phosphosite accessibility along their activation landscapes. Our results provide a systematic framework for comparative kinase ensemble characterization, integrating DFG geometry with A-loop and other regulatory elements to resolve kinase-specific states and the atomic-resolution structural rearrangements underlying their interconversion.

## 1 Introduction

Protein kinases are central regulators of eukaryotic cellular signalling. More than 500 protein kinases are encoded in the human genome, accounting for approximately 2% of human genes (1). Kinases control diverse processes including cell growth, proliferation, differentiation, and apoptosis through phosphorylation of protein substrates (2; 3). Given their central role in all aspects of cell life, kinase activity is tightly regulated through mechanisms such as autoinhibition, phosphorylation, and allosteric modulation (4; 5; 6). Disruption of their activity by mutation or dysregulated expression is often associated with many human diseases, such as a variety of cancers, emphasizing kinases as one of the most important target classes in drug discovery (7; 8; 9; 10).

Kinase activity is largely rooted in its catalytic kinase domain (KD), a highly conserved domain across the protein kinase family that mediates phosphorylation activity through conformational changes. KDs have a bilobal structure, consisting of a smaller N-terminal lobe and a larger C-terminal lobe, with the ATP-binding site located in the cleft between them (Supplementary Fig. S2). Several conserved structural motifs have been found to play central roles in kinase catalysis and regulation (11). Briefly, the glycine-rich loop (G-loop) in the N-lobe helps position and stabilize ATP, and the *β* 3 strand contains a conserved lysine that forms a salt bridge with the *α*C helix to organize the active-site architecture. In the C-lobe, the activation loop (A-loop) is a flexible segment that controls substrate access (12). At the N-terminus of the A-loop lies the highly conserved Asp-Phe-Gly (DFG) motif, whose Asp residue coordinates a divalent magnesium ion that helps position the ATP phosphates for catalysis (11; 13; 14; 15). Together, conserved elements organize the catalytic machinery required for phosphate transfer, and their coordinated rearrangements provide a structural basis by which kinases switch between functionally distinct conformational states (4; 11; 15).

Kinase activity is not determined by a single static structure but by the dynamic equilibrium between active and inactive states. States are commonly distinguished by characteristic arrangements of the DFG motif, the *α*C helix, and the A-loop (Supplementary Fig. S2). Modi and Dunbrack have determined that three main conformational states—DFG-in, DFG-out, and DFG-inter—can be defined according to two distances between these structural elements (*D*_1_ and *D*_2_, Fig. 1b) (14; 16; 17). They further noted how these main states can be subdivided according to specific values of *φ*, *ψ* and *χ*_1_ angles within the DFG motif. The DFG-in BLAminus state corresponds most closely to the active kinase conformation where catalytic metal ions bind (see Methods) (14). However, the DFG-motif does not fully define the functional conformational states. The A-loop that extends from the DFG-motif and controls substrate access has shown to undergo substantial rearrangement between the active and inactive functional states (18; 19; 20). In the active state, the A-loop adopts an extended, open conformation compatible with substrate binding, whereas inactive states can involve partial or complete folding of the A-loop over the substrate-binding region. DFG-out conformations can occur with either an active-like A-loop or a substantially reorganized folded A-loop, giving rise to structurally distinct inactive states (14).

**Figure 1:**
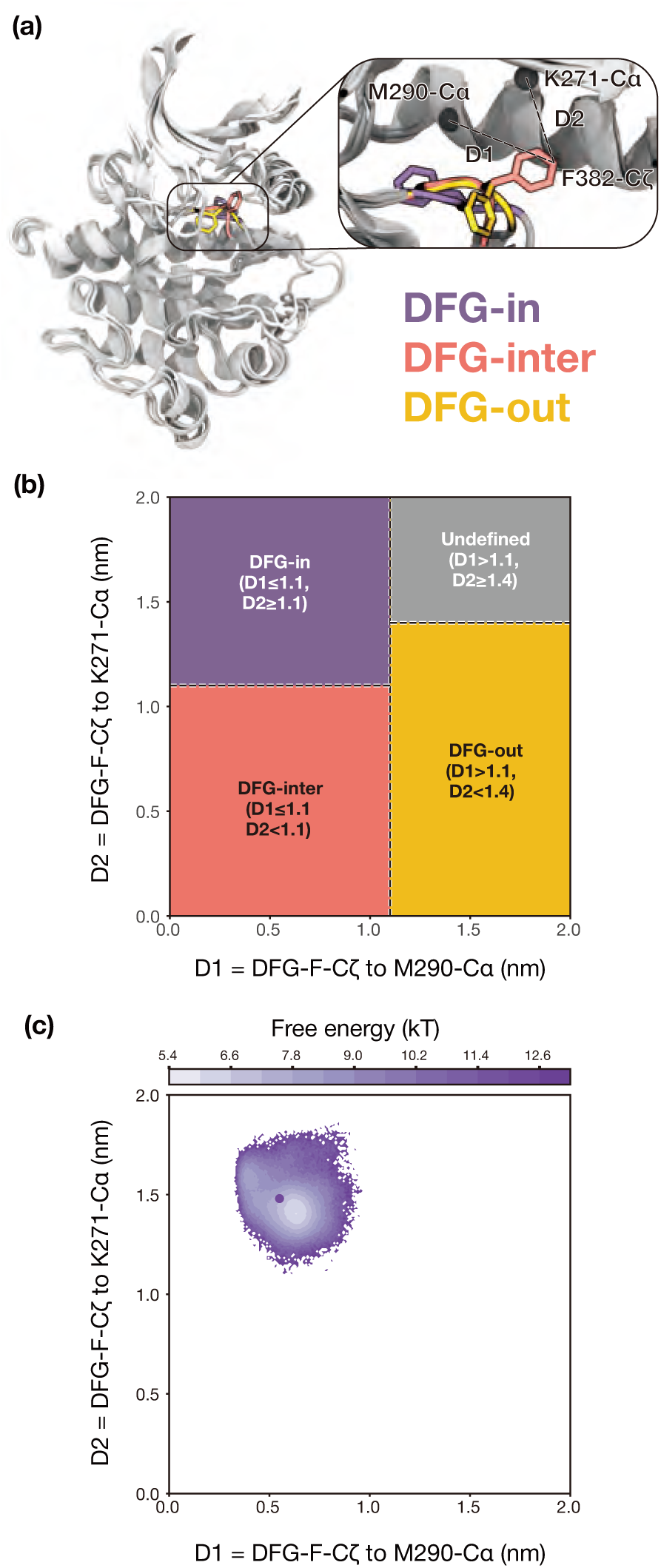
DFG motif conformational states of kinase domains. **(a)** Structural overlay of human ABL kinase in three conformations: DFG-in (PDB: 2F4J, purple), DFG-inter (PDB: 4XEY, salmon), and DFG-out (PDB: 1OPL, yellow). Inset: zoomed-in view of the DFG motif. D_1_ and D_2_ refer to distances between *α*C helix-M290-C*α* and DFG-F382-C*ζ*, and between *β* 3-K271-C*α* and DFG-F382-C*ζ*, respectively. **(b)** Empirical partitioning of DFG states based on D_1_ and D_2_. **(c)** Aggregate 1235.39 *µ*s simulation initiated from a DFG-in conformation (PDB: 2WGJ, purple point) projected onto D_1_ and D_2_, showing confinement to the DFG-in basin.

Ultimately, kinase activity is determined by the relative populations of these aforementioned conformational states. (4; 17; 21; 22; 23). Changes in the active–inactive equilibrium are central to normal regulation, for example, through phosphorylation or ligand binding, but can also contribute to diseases. Indeed, several clinically observed kinase mutations bias the conformational ensemble towards the active state, resulting in dysregulated signalling and oncogenic activation (24; 25; 26). Conversely, kinase inhibitors exploit these conformational preferences through distinct binding modes (10; 27). Type I inhibitors bind competitively in the ATP pocket, whereas type II inhibitors extend into an adjacent hydrophobic back pocket that is typically exposed in the DFG-out inactive conformation. By stabilizing inactive states, type II inhibitors are broadly observed to achieve improved selectivity (28; 29). Allosteric inhibitors bind outside the ATP pocket and can stabilize distinct inactive or autoinhibited conformations, offering opportunities for high specificity and reduced resistance (30; 31). Thus, understanding how genetic variants, post-translational modifications, and ligand binding modulate kinase function requires a quantification of how these events and mutations modulate kinase structural ensembles (25; 32; 33; 34; 35). Previous efforts have highlighted that some protein families follow common activation mechanisms, although it remains unclear if tyrosine KDs share a common activation ensemble (36). An ensemble-level characterization expands opportunities for structure-based drug design. The discovery of cryptic or transient states opens new opportunities for selective targeting to modulate kinase activity and potentially overcome resistance (37; 38; 39; 40; 41).

Despite their importance, kinase conformational ensembles remain challenging to characterize. Experimental techniques such as X-ray crystallography and cryo-EM provide high-resolution structural snapshots but typically capture static or averaged structures and commonly observe the most populated conformational states (42). Solution methods such as NMR and hydrogen–deuterium exchange can report on conformational dynamics but often provide limited atomistic detail for low-population intermediates (43). Molecular dynamics (MD) simulations provide a complementary route to probe kinase dynamics at atomic resolution, yet functionally important transitions, such as DFG flips and coupled A-loop or *α*C helix rearrangements, can occur on millisecond timescales and may remain inaccessible to conventional equilibrium simulations initiated from a single starting structure (15; 20; 44; 45; 46). Consistently, our own single-starting-point unbiased simulations struggle to sample the ensemble (Fig. 1c), showing that even 1.24 ms of unbiased simulations initiated from an experimentally determined DFG-in conformation remains trapped in this state. Enhanced-sampling methods can accelerate barrier crossing and discover rare states, but quantitative interpretation of the resulting ensembles can depend on the chosen collective variables, biasing protocol, and reweighting procedure (39; 47; 48; 49; 50). As a result, no singular protocol has observed multiple activation transitions across many KDs, and it remains unclear whether the activation pathway follows a conserved mechanism across kinases.

Markov state models (MSMs) provide a framework to address this challenge. By integrating information from multiple unbiased MD simulations, MSM analysis enables the identification of stable states and estimating transition timescales between them. Kinases are well-suited to this approach since many experimental structures spanning active and inactive states are available (12), providing seeds for sampling relevant conformational states (39; 51; 52; 53). However, for structurally under-represented kinases, experimental structures may not cover all relevant functional states. AI-based structure predictors offer a collective-variable-independent way to generate diverse structural seeds without first carrying out expensive exploratory simulations. In particular, AlphaFold2 (AF2) can be guided by template structures, enabling structural information from well-characterized homologues to be transferred onto a target sequence (54; 55; 56). For conserved protein families such as kinases and GPCRs, homologous structures annotated by conformational state can be used to generate physically plausible starting models in states that are poorly represented in the target’s experimental structural ensemble (46; 56; 57; 58). We refer to this strategy as transfer seeding, where homologue-derived conformational templates generate key target-specific seeds for subsequent unbiased MD and MSM analysis.

Here, we characterized the equilibrium conformational landscapes and activation pathways of three apo tyrosine KDs, ABL1 (which we refer to as ABL), EGFR, and MET, using MSMs constructed from unbiased MD simulations. We generated large aggregate unbiased simulation datasets comprising 0.592 ms for ABL, 1.836 ms for EGFR using PDB seeds, and 5.938 ms for MET on the Folding@home distributed computing platform (41) (Supplementary Fig. S6). For MET, we introduced the transfer-seeding protocol to complement experimental structures with AF2-generated conformations, expanding the initial structural diversity available for unbiased sampling. By sampling each kinase with a consistent, unbiased simulation and MSM analysis protocol, our simulations enable direct comparison of their conformational ensembles. Although the three kinases sample similar active-like and inactive conformational states, we find that their equilibrium populations differ substantially. The three kinase activation pathways also proceed through distinct intermediate states and interconvert on timescales ranging from approximately 100 *µ*s to 6 ms. We further find that the A-loops define kinase-specific conformational ensembles that shape the transition pathways. Our work establishes a systematic biophysical framework for comparing kinase activation landscapes, enabling mechanistic discovery of how conserved structural elements give rise to kinase-specific conformational dynamics.

## 2 Results

### 2.1 ABL and EGFR share common DFG conformational states but differ in populations and kinetics

We sampled kinase conformational ensembles using a diverse set of PDB seeds for two well-represented KDs ABL and EGFR (Supplementary Table S1, Fig. S4, S5). Projection onto the *D*_1_ and *D*_2_ distances showed broad coverage of the DFG-in, DFG-inter, and DFG-out regions (Fig. 2a, Fig. S4, S5). For MSM construction, we used a feature set that captures the rearrangement of the DFG-motif, αC helix, and A-loop (see Methods) and identified six macrostates for both ABL and EGFR (Fig. 2b). To relate these macrostates to experimentally observed kinase conformations, we labelled them with the kinome-wide DFG motif classification introduced by Modi and Dunbrack (14) (see Methods) and characterized their A-loop conformational ensembles.

**Figure 2:**
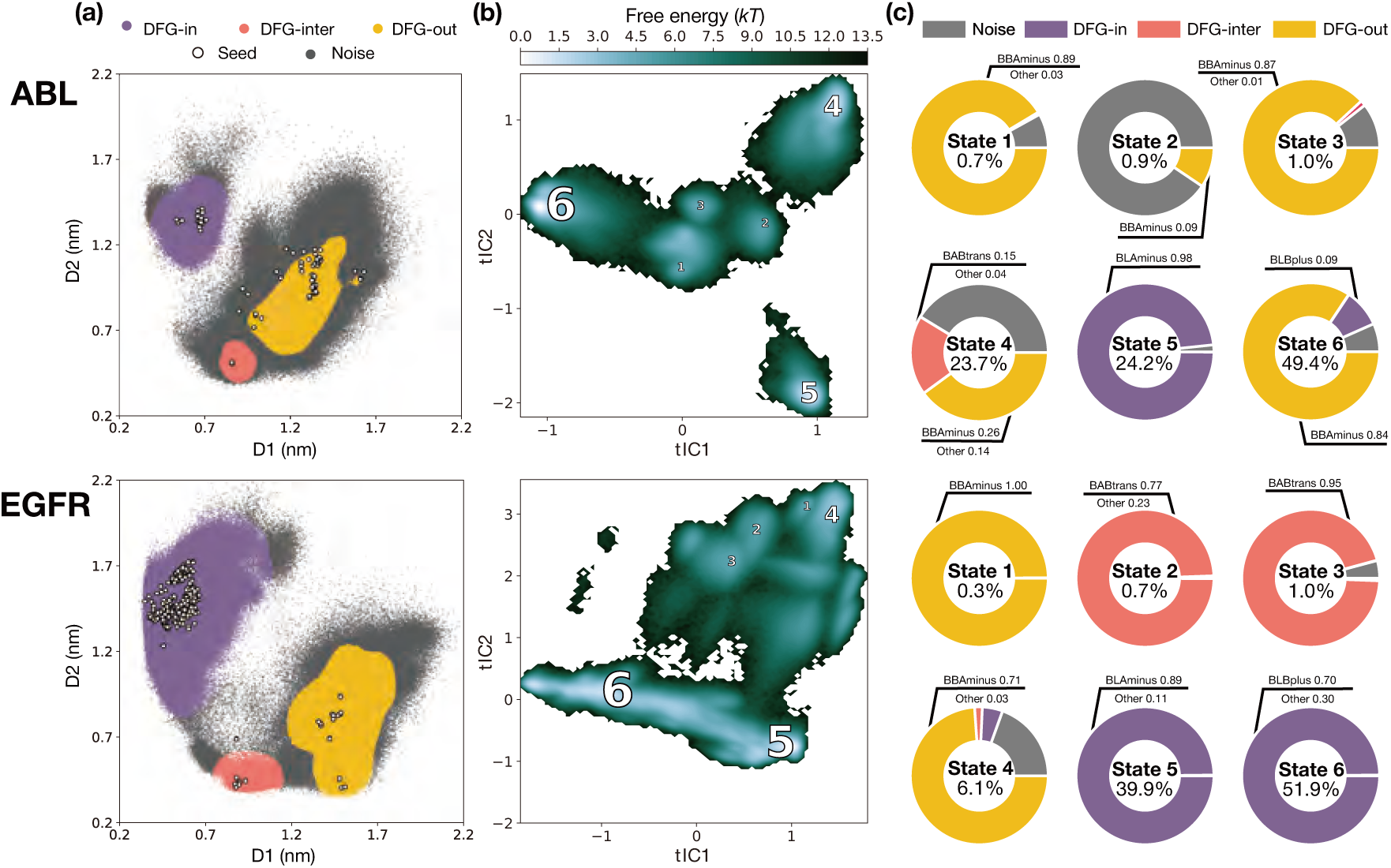
MSMs of ABL and EGFR explore similar DFG states but differ in population and landscapes. **(a)** DFG-state classification of MD simulation frames based on *D*_1_ and *D*_2_ distances. Each MD frame is a points coloured by assignment to DFG-in (purple), DFG-inter (salmon), DFG-out (yellow), or undefined (gray). Initial seeding structures are marked as white circles. **(b)** ABL and EGFR free energy landscapes estimated from MSM equilibrium distribution projected onto tIC1 and tIC2. Macrostates are labelled with sizes proportional to their populations. **(c)** Pie charts show the decomposition of each macrostate into Dunbrack states. Labels report macrostate equilibrium population and Dunbrack state fractions.

Our MSMs show that ABL and EGFR access a shared subset of DFG conformational states, but assign different equilibrium populations to them. These include the active-like DFG-in BLAminus state, the SRC-like inactive DFG-in BLBplus state (16; 59), the DFG-out BBAminus state, and the DFG-inter BABtrans state (Fig. 2c). Ensemble images of each macrostate labelled with DFG, αC helix, and A-loop states are provided in Supplementary Figs. S11 and S12. In their respective six-state coarse-grained MSMs, most macrostates are dominated by a single Dunbrack class (Fig. 2c), indicating that the kinome-wide DFG conformation nomenclature captures a substantial part of the slow conformational heterogeneity resolved by the simulations. In apo ABL, the equilibrium ensemble is dominated by macrostate 6 (49.4%). It is primarily composed of DFG-out BBAminus, and A-loop closed conformation. This arrangement resembles a typical inactive state stabilized by type II inhibitors with PDB 6XRG being an example. The second most populated ABL macrostate, state 5 (24.2%), corresponds to the DFG-in BLAminus and A-loop open ensemble, which is characteristic of catalytically active-like kinase structures (example PDB 2F4J) (17). The EGFR equilibrium ensemble is dominated by DFG-in conformations. State 6 (51.9%) corresponds to the BLBplus ensemble and adopts a SRC-like inactive conformation, in which the DFG-Asp remains oriented toward the ATP-binding site, the regulatory *β* 3-Lys–αC-Glu salt bridge is disrupted, and the A-loop partially folds into a two-turn helix that interacts with the displaced αC helix (Supplementary Fig. S19). The second most populated EGFR macrostate 5 (39.9%) corresponds to the active-like BLAminus ensemble with the A-loop stays open. We also identify state 1 as under-represented in the experimental structure set, characterized by an untypical open A-loop with closer packing between its central and C-terminal segments, while retaining an overall inactive conformation.

The correspondence between DFG labels and MSM macrostates is strong but not absolute. DFG classifications describe local structural geometries of the motif, but do not necessarily imply kinetic similarity or separation between conformational states. We quantified the interconversion timescales between macrostates using mean first-passage times (MFPTs), defined as the average time required for trajectories starting in one state to reach a target state for the first time (Fig. S16). The ABL MSM macrostate 6 includes an SRC-like inactive substate in the BBAminus ensemble (Supplementary Fig. S17). The MSM shows the SRC-like inactive substate interconvert rapidly with the classical inactive conformations, with an MFPT of 68.2 *±* 0.20 *µ*s, whereas the transition from macrostate 6 to the active-like BLAminus macrostate 5 occurs on a much slower timescale of 2.39 *±* 0.20 ms. The DFG-out BBAminus-to-DFG-in BLBplus transition is faster than the subsequent BLBplus-to-BLAminus rearrangement within the DFG-in ensemble, showing that changes in broad DFG classification do not directly reflect kinetic barriers for ABL. A similar decoupling between DFG classification and kinetic state identity is observed in ABL macrostate 4, which contains kinetically mixed DFG-out BBAminus and DFG-inter BABtrans conformations (Supplementary Fig. S18). These conformations share a similar open, minimally perturbed A-loop ensemble and differ primarily in the local dihedral geometry of the DFG motif and the intermittent formation of the *β* 3-Lys–αC-Glu salt bridge. A related experimental example is provided by PDB 2G2F, whose two ABL chains retain similar minimally perturbed A-loop conformations while adopting distinct DFG geometries, illustrating that substantial rearrangement of the DFG motif can occur without a corresponding large-scale reorganization of the A-loop (60; 61; 62). These observations show that DFG labels provide a useful structural annotation, but they do not define kinetically distinct states or predict their connectivity.

### 2.2 Transfer seeding expands coverage of the MET kinase DFG-state ensemble

MET is less represented in the experimental structural record (Fig. 3c, S20). There are no MET DFG-inter structures available in the PDB, leaving a gap between the experimentally observed DFG-in and DFG-out regions (Fig. 3a, S20). Due to the rarity of DFG transitions on conventional MD timescales, simulations initiated only from this limited PDB ensemble are unlikely to identify missing intermediate states de novo.

**Figure 3:**
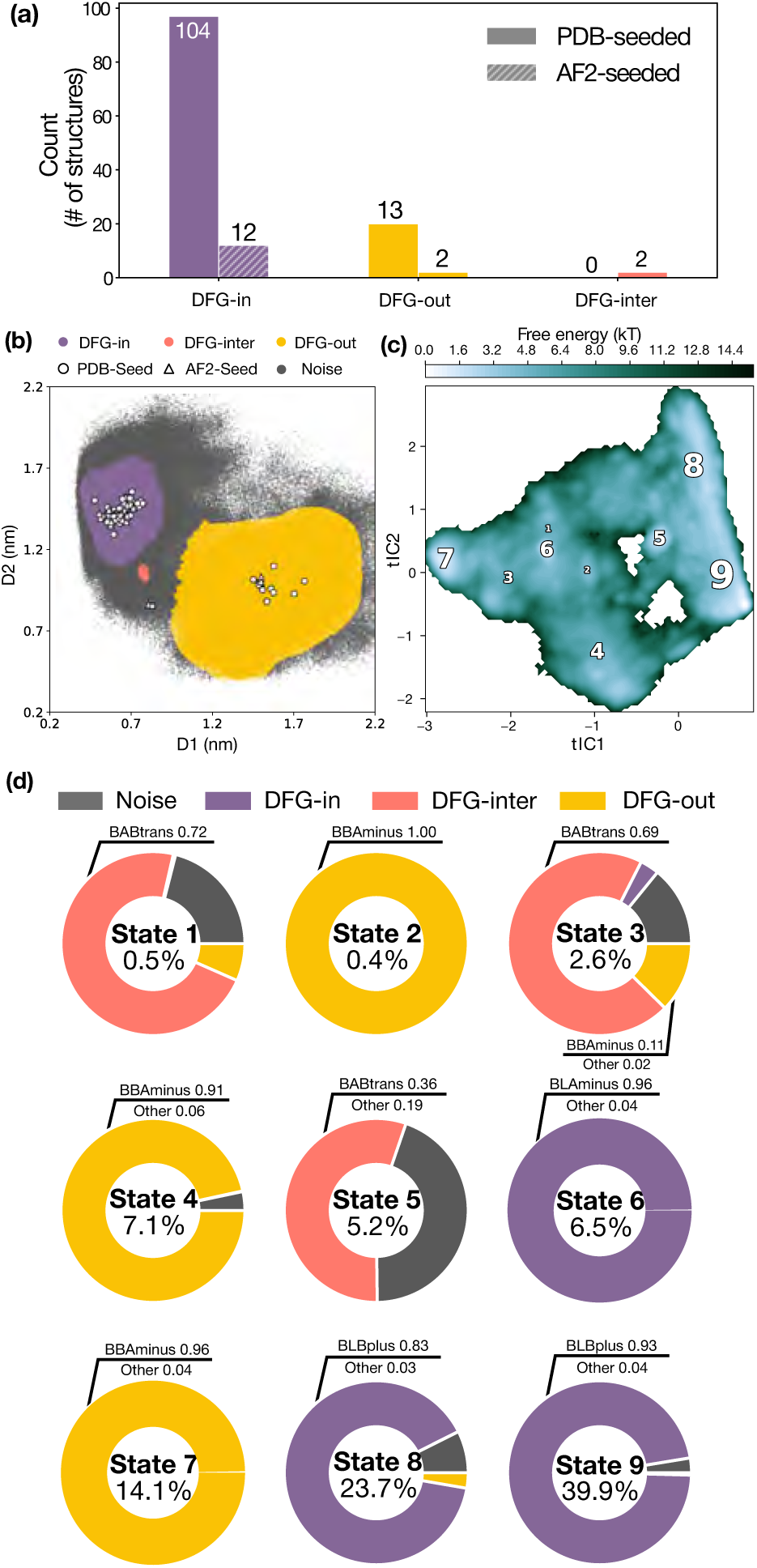
Complete sampling and MSM construction for MET kinase are enabled by transfer seeding. **(a)** Number of MET PDB structures (solid) and AF2-derived seeds (hatched) used for simulations. **(b)** Simulation snapshots projected onto *D*_1_ and *D*_2_ distances, coloured by DFG-state assignment as in Fig. 2. Initial seeding structures are marked to indicate PDB seeds (white circles) and AF-derived seeds (white triangles). **(c)** MET free energy landscape estimated from MSM equilibrium distribution projected onto tIC_1_ and tIC_2_. Macrostates are labelled with sizes proportional to their populations. **(d)** Pie charts show the decomposition of each macrostate into Dunbrack states. Labels report macrostate equilibrium population and Dunbrack state fractions.

We first tested whether missing MET DFG states could be recovered from the DFG-in basin using enhanced sampling approaches, including accelerated MD, Gaussian accelerated MD, umbrella sampling, metadynamics, and adaptive sampling (see Methods and SI). All enhanced sampling approaches used the same CVs, geometric distances *D*_1_ and *D*_2_, to ensure that the biases used were not CV-insensitive. Several of these approaches recovered DFG-inter and DFG-out conformations, indicating that the experimentally missing DFG-inter region is physically accessible. However, the apparent free-energy landscapes differed substantially between protocols, particularly in the relative depths of DFG basins and transition regions (Supplementary Fig. S1). We attribute this to a combination of biasing and reweighting uncertainty, incomplete barrier crossing, and limitations of low-dimensional collective variables. Meanwhile, some conformational changes sampled in unbiased MD, as seen in ABL and EGFR, involve coupled rearrangements and are not cleanly described as motion along *D*_1_ and *D*_2_ alone. Therefore, these biased simulations, without an in-depth consideration of multiple viable collective variables (57), are mainly for state-discovery, but are insufficient for quantitative estimates of the equilibrium ensemble. To systematically generate a MET ensemble in a similar manner to ABL and EGFR without introducing biasing potentials, we adopted a parallel simulation strategy initialized from diverse conformations (39). However, the available experimental MET structures do not cover all relevant DFG conformational states, limiting the diversity of initial simulation seeds. AF2 modelling of the MET sequence with default settings and inputs also failed to recover DFG conformations other than DFG-in (Supplementary Fig. S3). To overcome these limitations, we developed the transfer-seeding approach in which annotated kinase structures from KinCore were provided as conformational templates to AF2, transferring structural information from ABL and EGFR onto the MET sequence (see Methods). This approach successfully generated MET structures in Dunbrack states absent from the available MET PDB ensemble and not recovered by untargeted AF2 modelling (Supplementary Fig. S14). AF2 was found more reliable at generating the desired templated conformations than the other co-folding methods evaluated (Supplementary Fig. S15). To expand the initial conformational diversity of MET while limiting the contribution from predicted structures, we include two AF2-generated structures per Dunbrack state (Fig. 3a).

Our simulations identify MET conformational ensembles that are not represented in the available experimental structural set. The union of PDB- and AF2-seeded simulations populates a connected landscape spanning DFG-in, DFG-inter, and DFG-out conformations (Fig. 3). The MSM resolves DFG-inter ensembles in states 1, 3, and 5, despite the absence of experimentally determined DFG-inter MET conformations, indicating that their absence from the PDB reflects incomplete structural coverage rather than an inaccessible region of the MET landscape (Fig. 3c,d). Ensemble representations of each macrostate, annotated by DFG, αC-helix, and A-loop conformations, are provided in Supplementary Fig. S13. States 1 and 3 represent low-population metastable DFG-inter intermediates in which the A-loop is partially folded and interacts with the αC helix while the helix remains in an inward orientation. The simulations additionally resolve an active-like BLAminus ensemble in state 6 (6.5%), characterized by an open A-loop and active-site geometry, while the existing active-like experimental MET structures predominantly adopt an ABAminus DFG geometry.

As in ABL and EGFR, MET conformational heterogeneity extends beyond the DFG motif, with A-loop rearrangements contributing substantially to the separation of major kinetic basins. States dominated by the same DFG label are not necessarily structurally or kinetically equivalent (Fig. 3c,d). The DFG-in ensemble separates into three distinct kinetic basins. State 6 (6.5%) corresponds to the active-like BLAminus ensemble with an open A-loop, whereas states 9 (39.9%) and 8 (23.7%) both adopt SRC-like inactive conformations characterized by a BLBplus DFG motif and a displaced αC helix. States 8 and 9 are distinguished by the C-terminal A-loop ensemble. Secondary-structure propensities show that this region remains predominantly coil-like in state 8, whereas state 9 exhibits increased *β* -strand propensity (Supplementary Fig. S23). Representative experimental structures are PDB 2G15 for state 8 and PDB 7B3W for state 9. Despite forming distinct metastable basins, states 8 and 9 interconvert rapidly, with MFPTs of 12.0 *±* 0.6 *µ*s and 19.0 *±* 0.7 *µ*s, respectively, indicating local A-loop exchange within the broader SRC-like inactive ensemble. In contrast, transitions from either state to the active-like state 6 require approximately 5.8 ms, whereas escape from state 6 back to states 8 or 9 occurs within 0.4–0.45 ms. This shows the BLAminus active-like basin is less accessible than the dominant SRC-like ensemble in the apo form. A similar subdivision occurs within the DFG-out ensemble. States 2, 4, and 7 are kinetically distinct despite sharing a BBAminus DFG geometry and a displaced αC helix, with their folded A-loops adopting distinct secondary-structure arrangements. Our results show that available experimental MET structures capture only a subset of its metastable conformational landscape, whereas transfer seeding and MSM analysis resolves additional metastable and intermediate conformations. When access from SRC-like BLBplus to active-like BLAminus conformations is compared across kinases, the transition is millisecond-slow in MET and ABL but occurs on a much shorter tens-of-microseconds timescale in EGFR, demonstrating that similar DFG-in structural classes can have different kinetic connectivity across KDs.

### 2.3 Activation transitions follow kinase-specific pathways

While we simulate the apo forms of the KDs, previous studies have shown that MD can capture key conformational intermediates along activation pathways that are accessible under thermal equilibrium (39; 40; 46; 63). We compute and compare activation pathways and intermediates for the ABL, EGFR, and MET KDs. We used transition path theory (64) to decompose reactive trajectories connecting a DFG-out (BBAminus) macrostate to a DFG-in (BLAminus) macrostate into reactive fluxes, representing the probability flow between the designated start and end states across MSM microstates (Fig. 4). For ABL and MET, the DFG-out endpoint was chosen from macrostates with a closed activation loop (A-loop), corresponding to the canonical inactive conformation. For EGFR, an analogous closed A-loop inactive state was not observed. We defined the EGFR DFG-out endpoint using a BBAminus macrostate with an open A-loop and an outward-displaced *α*C helix. For each KD, we report the highest-flux pathway and define its rate-limiting step as the transition with the smallest flux. Transition path ensembles are provided in Supplementary Fig. S24. To systematically identify structural changes accompanying the rate-limiting step, we trained Random Forest classifiers to distinguish pre-versus post-rate-limiting conformations and extracted the most discriminating structural features (see Methods).

**Figure 4:**
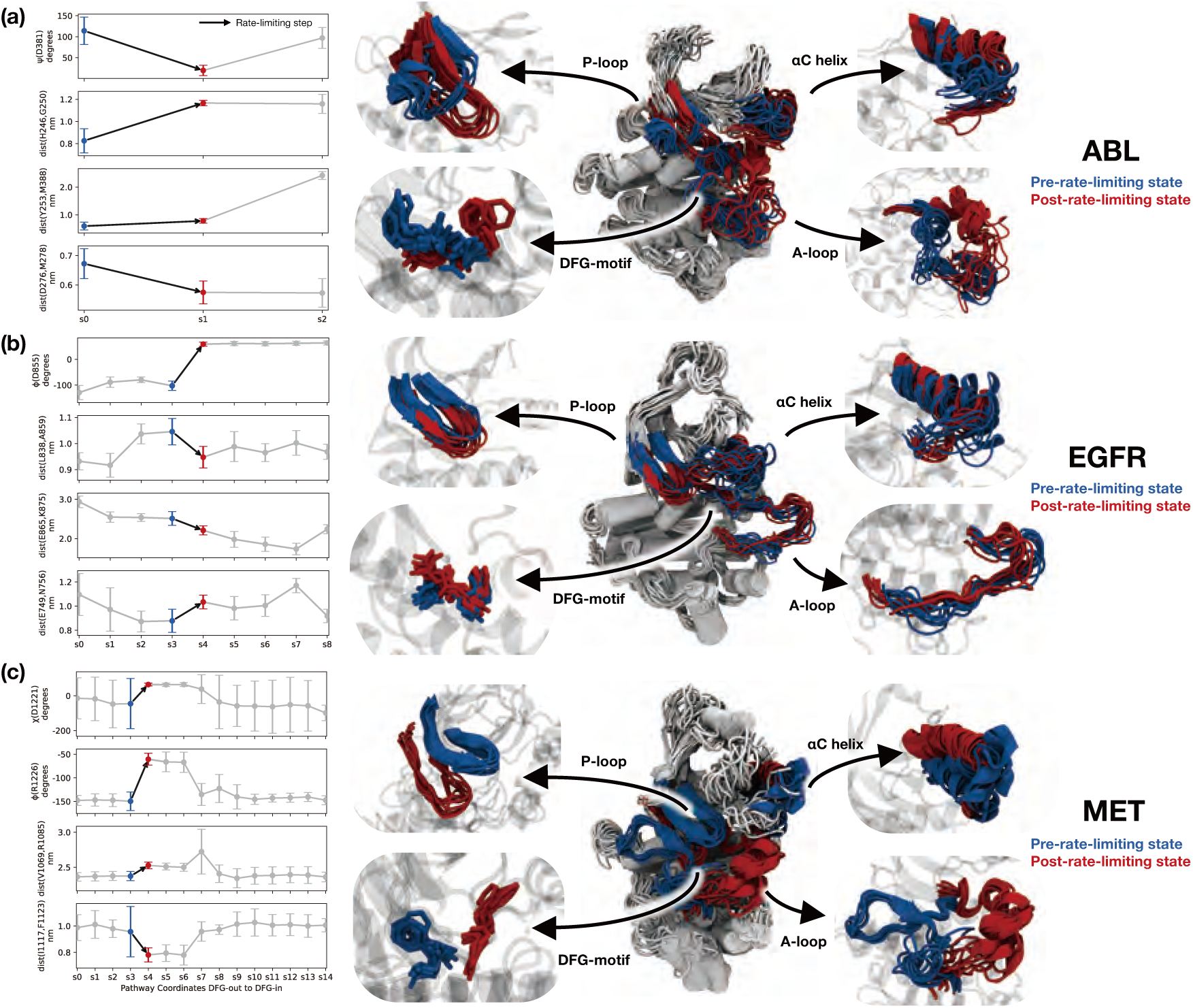
Rate-limiting steps in the DFG-out to DFG-in transition. Analysis for ABL **(a)**, EGFR **(b)**, and MET **(c)**. Left: mean and standard deviation of top-ranked features from the random forest classifier along the transition pathway. The rate-limiting step is indicated by black arrows; pre- and post-rate-limiting states are shown as blue and red markers respectively. Right: superposition of pre-rate-limiting (blue) and post-rate-limiting (red) conformations, overlaid on the full ensemble (grey). Insets highlight key regulatory elements.

The ABL highest-flux pathway contains two major transitions. The first transition, from the inactive BBAminus state with a closed A-loop to the SRC-like inactive ensemble, forms the principal flux bottleneck (Fig. 4a). This rearrangement is initiated by outward displacement of the αC helix, which disrupts the *β* 3-Lys–αC-Glu salt bridge and opens the back pocket. The resulting increase in space allows the A-loop to rotate towards a SRC-like orientation and form a two-turn helix packed against the αC helix. Concurrently, the G-loop rearranges from a short helical conformation into a more extended turn projecting towards the ATP-binding site. In the subsequent transition from the SRC-like inactive intermediate to the active-like BLAminus ensemble, the A-loop rotates further into the open conformation, while the αC helix moves inward to re-form the *β* 3-Lys–αC-Glu salt bridge and the G-loop returns to a kinked conformation.

The EGFR highest-flux pathway is dominated by comparatively local rearrangements of the DFG geometry, rather than the larger-scale A-loop and αC-helix motions observed for ABL (Fig. 4b, Supplementary Fig. S24b). The pathway begins with rearrangement of residue Ala854 preceding the DFG motif, converting the DFG-out BBAminus state into an inactive DFG-in ABAminus intermediate. The subsequent ABAminus-to-BLAminus transition is the rate-limiting step and is characterized by reorganization of the Ala854-DFG backbone dihedrals, accompanied by subtle rearrangement of the A-loop, which remains open throughout the transition. Minimal structural changes are observed in the G-loop or αC helix during this rate-limiting step. Notably, the SRC-like BLBplus ensemble does not participate in the dominant net-flux pathways, indicating that it forms an off-pathway basin that predominantly interconverts with the active-like BLAminus ensemble.

In MET, the highest-flux pathway resolves a progressive opening of the A-loop coupled to coordinated rearrangements of the DFG motif, αC helix, and G-loop, proceeding from the BBAminus ensemble (state 7) through two BABtrans intermediates (states 1 and 3) before reaching the BLAminus state 6 (Fig. 4c, Supplementary Fig. S24c). Notably, states 1 and 3 are intermediate conformations not represented in the experimental structural set and are instead resolved through simulations. Along this pathway, the A-loop progressively approaches the active-like conformation, with its RMSD to the active-like state decreasing from 21.9 Å in state 7 to 11.4 Å in state 1 and 3.6 Å in state 3. The rate-limiting transition from state 7 to state 1 is associated with the initial opening of the A-loop, which forms helical elements that interact with the αC helix while the DFG motif adopts the BABtrans geometry. This transition is accompanied by formation of the *β* 3-Lys–αC-Glu salt bridge and extension of the G-loop into the space released by A-loop opening. The subsequent transition from state 1 to state 3 involves further opening of the A-loop and dissociation of these helical elements, after which the final transition from state 3 to state 6 completes the BABtrans-to-BLAminus rearrangement. The path ensemble also contains lower-flux branches connecting both states 1 and 3 directly to the sink state 6, indicating that although the described route carries the highest flux, more direct routes to the active-like BLAminus ensemble are also accessible (Supplementary Fig. S24c).

### 2.4 Activation loop phosphosite exposure precedes activation transition in distinct conformational intermediates

The A-loop is a dynamic regulatory element in kinases. Its flexibility enables conformational changes that regulate substrate access, but also means that the A-loop is frequently unresolved or partially resolved in experimental crystal structures (65). A-loop phosphorylation provides a common regulatory switch by stabilizing catalytically competent conformations (66), although the conformational states in which phosphorylation sites (phosphosites) become solvent-accessible can differ between kinases. To examine coupling between phosphorylation site accessibility and A-loop dynamics in our MSMs, we characterized the A-loop ensemble of each macrostate using per-residue secondary-structure labels assigned by DSSP (see Methods) and relative solvent-accessible surface area (rSASA), normalized from 0 to 1 for each residue.

In the active-like BLAminus ensembles, the phosphosites ABL Tyr393 (66) and MET Tyr1235 (67) occupy similarly packed local environments. DSSP analysis shows that both residues lie within locally structured A-loop segments containing *β* -strand and turn conformations, rather than within fully disordered loops (Supplementary Figs. S21 and S23). In both ABL and MET, the phosphosite side chains are partially shielded by neighbouring bulky or aliphatic residues, resulting in relatively low solvent exposure in the active-like BLAminus ensemble (Fig. 5a,c).

**Figure 5:**
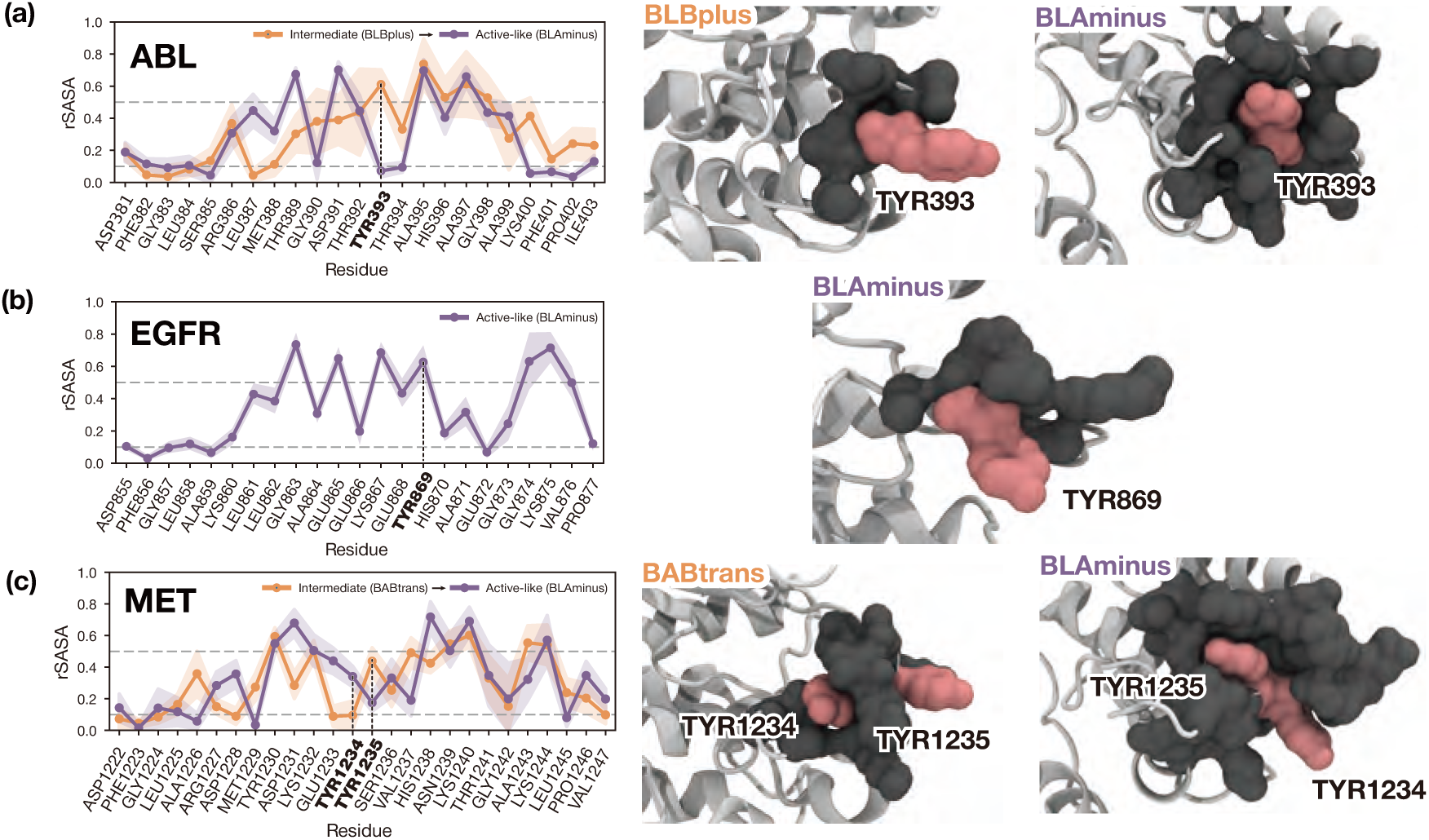
A-loop phosphorylation-site exposure is kinase specific, with Tyr393 (ABL) and Tyr1235 (MET) being more accessible in intermediate ensembles than in the active-like ensemble. Analysis for ABL **(a)**, EGFR **(b)**, and MET **(c)**. rSASA profiles (left column) and representative A-loop conformations (right column) are shown for the active-like ensemble (colored purple, right) and, where active-state (purple) phosphosites are buried, a more exposed intermediate ensemble (colored orange, right). Lines show mean rSASA and shaded regions indicate standard deviation for the sampled conformations; horizontal dashed lines mark rSASA values of 0.1 and 0.5 to indicate complete burial or complete solvent exposure; vertical dashed lines mark phosphosite residues. Phosphosite residues are shown as pink surfaces in the structure. Surrounding residues are shown in grey surfaces.

This state-dependent exposure suggests that the apo active-like ensemble may not be the main phosphorylation-competent conformation of KDs. For MET, this interpretation is consistent with experimental evidence that A-loop phosphorylation is sequential, with Tyr1235 phosphorylated before Tyr1234 (68; 69). In the highest-flux pathway, MET activation proceeds from the BBAminus inactive state through an intermediate BABtrans state before reaching the active-like BLAminus state. In the intermediate, Tyr1235 is more exposed (rSASA 0.44 *±* 0.09), whereas Tyr1234 remains largely buried (rSASA 0.10 *±* 0.09). Tyr1234 becomes more exposed only after progression towards the active-like BLAminus state (rSASA 0.35 *±* 0.06). These observations suggest that the BABtrans ensemble may represent a pre-active, phosphorylation-competent intermediate for the first MET A-loop phosphorylation event at Tyr1235, which promotes subsequent phosphorylation at Tyr1234 and A-loop rearrangement towards the active-like ensemble.

The same principle of state-dependent phosphosite accessibility might apply to ABL. Tyr393 is less solvent-exposed in the active-like BLAminus state (rSASA 0.07 *±* 0.04) than in the intermediate BLBplus ensemble (rSASA 0.61 *±* 0.09), where Tyr393 points towards solvent (Fig. 5a). This ensemble is also involved in ABL highest-flux pathway as the post-rate-limiting step. We therefore hypothesize that ABL Tyr393 phosphorylation is preferentially enabled by a BLBplus pre-active intermediate that presents Tyr393 in a more solvent-accessible geometry, rather than by the final apo active-like BLAminus state.

EGFR presents a contrasting picture. In the active-like BLAminus ensemble, the A-loop remains more open and predominantly coil-like around Tyr869. The Tyr869 side chain points outward towards solvent and is therefore highly exposed (Fig. 5b). This is consistent with EGFR’s non-canonical regulatory mechanism, in which catalytic activation is primarily mediated by asymmetric kinase-domain dimerisation rather than by a required A-loop phosphorylation switch (70; 71). In this context, Tyr869 exposure may reflect regulatory accessibility for cross-talk phosphorylation rather than a prerequisite step for forming the active catalytic state.

Given these observations from pre-phosphorylation apo ensembles, we hypothesize that A-loop phosphorylation sites are exposed in kinase-specific conformational windows along the activation pathway such that phosphorylation may select or stabilize transient pre-active ensembles rather than marking the final active state. This hypothesis may be tested by combining conformation-selective perturbations with HDX-MS (72), NMR (73; 74), or time-resolved phosphosite mass spectrometry (75) to determine whether phosphosite exposure and phosphorylation order track enrichment of the predicted intermediates.

## 3 Conclusions

Our simulations and MSMs show that kinase activation does not proceed through a conserved mechanism shared across tyrosine kinases, but through kinase-specific transitions embedded within broader regulatory ensembles. A key component of this comparison is the transfer-seeding strategy, which expands structurally under-represented starting ensembles using homologue-guided AF2 models and enables their subsequent characterization with the same unbiased MD and MSM framework used for well-represented kinases. Although ABL, EGFR and MET can access a common subset of DFG geometries, their equilibrium populations, kinetic connectivity and dominant activation routes differ substantially. The simulations resolve metastable conformations not represented in the available experimental structural set, including low-population intermediates along the MET DFG-activation pathway and additional A-loop arrangements in EGFR and MET. Building on established DFG structural classifications, including the Dunbrack nomenclature, our MSM framework places these conformations within kinase-specific kinetic networks and resolves how structurally related states are connected through distinct transition mechanisms. Consistent with its established regulatory role, the A-loop provides an additional level of conformational resolution, distinguishing metastable ensembles, shaping activation pathways, and modulating phosphosite exposure. Our results provide an ensemble-level description of kinase activation that resolves the coupled conformational diversity of the A-loop and other regulatory elements at atomic resolution, placing conformational states within a dynamic picture of kinase-specific activation landscapes. This mechanistic heterogeneity has direct implications for interpreting kinase regulation and for conformation-selective inhibitor design. The accessibility and stability of DFG-out and intermediate states, together with the coupling among the DFG motif, *α*C helix, G-loop, and A-loop, differ across kinases. Inhibitor selectivity may depend on the broader conformational network through which active and inactive states are populated and interconverted, rather than simply the local interaction of the inhibitor binding site (32). For example, the SRC-like ABL intermediate and the low-population MET intermediate with a partially open A-loop illustrate how kinase-specific conformations may provide opportunities for selectively stabilizing inactive or intermediate states.

Transfer seeding provides a practical route for extending this systematic ensemble-generation and MSM analysis to kinases with incomplete experimental structural coverage. For MET, homologue-guided AF2 models broaden the starting ensemble and enable sampling of connected DFG-inter and DFG-out regions that are not represented in the available experimental structures. This strategy complements adaptive and enhanced-sampling approaches by using conformational information already present in related family members to initialise unbiased simulations. The present models are limited to isolated apo KDs, while ligand binding, phosphorylation, mutations and regulatory domains may reshape both populations and pathways (33; 34; 35; 76; 77). We envision that extending this framework to perturbed ensembles and linking functional insights with ensemble and kinetic descriptions will open new avenues for the design of next-generation kinase inhibitors with improved selectivity and affinity.

## 4 Methods

### 4.1 Structural seed generation

For each target kinase domain (KD) we assembled a pool of starting conformations (“seeds”) intended to span the DFG-in, DFG-inter, and DFG-out regions of the Modi–Dunbrack *D*_1_ vs. *D*_2_ landscape (14). Seeds were drawn from two sources: (i) experimentally determined structures available in the Protein Data Bank (PDB-based seeding), and (ii) AlphaFold2 (AF2) models generated with ColabFold using curated template sets biased towards specific Dunbrack substates (AF2-based, or transfer seeding). All seeds were processed through an identical structure-preparation and solvation pipeline so that down-stream simulations differed only in their starting coordinates. Scripts for seed generation are available at https://github.com/sukritsingh/kinase-seeding.

### 4.2 PDB-based seed generation

Apo KD seeds for ABL1 (KLIFS kinase ID 392) and EGFR (KLIFS kinase ID 406) were generated from every structure available for the corresponding kinase in KLIFS (78). For each kinase, the full set of PDB entries, chains, and alternate models was queried with the opencadd KLIFS remote interface and filtered to retain only entries with reported resolution. Each (PDB ID, chain, alternate location) tuple was then passed as a PDBProtein component to the KinoML OEKLIFSKinaseApoFeaturizer (79), which uses the OpenEye Spruce toolkit with the RCSB loop database (rcsb_spruce.loop_db) to strip ligands, waters, and cofactors, model missing loops, repair missing side chains, assign protonation states at pH 7.4, add caps, and return an apo KD. Because the individual structures cover different residue ranges, all apo models for a given kinase were cropped to the common residue window (maximum of the per-structure minimum residue, minimum of the per-structure maximum residue), and any structure with internal gaps in that window was discarded; the retained structures were then re-capped with ACE/NME and residues were renumbered using assign_caps and update_residue_identifiers so that all seeds for a given kinase shared an identical topology. The resulting per-PDB apo structures constitute the PDB-based seeds: 83 seeds for ABL1 and 400 seeds for EGFR.

### 4.3 Transfer seeding of MET via AF2

For MET (KLIFS kinase ID 446, UniProt P08581), the PDB ensemble is dominated by DFG-in conformations. To augment this ensemble, we used homologue-guided AF2 modelling (transfer seeding) (54; 55; 80) with MSA subsampling and stochastic dropout enabled. MET KD sequences were modelled using (i) the default ColabFold MSA server (MMseqs2), (ii) stochastic dropout enabled at inference, (iii) num_recycles = 3, and (iv) two models per prediction, to generate a minimal set of AF2 seeds of MET kinase.

Custom template sets derived from the KinCore database (16), sorted by Dunbrack state labels, were used to bias each prediction towards a specific DFG state. The sorted template library is deposited at https://osf.io/spu2a/, and was created by downloading and re-sorting all structures into a directory tree keyed by those labels (one sub-directory per spatial DFG state, and within each, one sub-directory per Dunbrack substate). To bias AF2 towards a chosen state for MET, the templates directory was pointed at the corresponding labelled sub-directory, so that only structures carrying that Dunbrack label were available as templates. Separate AF2 runs were executed for each Dunbrack state template pool. As a minimal set to supplement existing PDB structures, two models from each Dunbrack state were created, giving a total of 16 AF2-based seeds, and were pooled with the native MET PDB structures. This pooled set of 117 PDB-based structures and 16 AF2-seed structures, a total of 133 structures, was then used as the initial MET kinase simulation seeds. The combined set of 133 seeds was processed through the same KinoML apo-kinase featurizer and common-residue cropping described above to yield a single, topology-consistent set of MET seeds. A sample demonstration of using the KinCore-sorted template library to do transfer seeding is available as a Colab notebook at https://colab.research.google.com/drive/1Dcb5ZouYJQfyq1np-TBmKJE8DOAbA9dU.

### 4.4 Molecular dynamics simulations

All seeds (PDB-based for ABL1/EGFR; PDB- and AF2-based for MET) were parameterized and equilibrated with an identical OpenMM 8.0 protocol (81), using AMBER ff14SB for protein (82) and TIP3P for water. Each apo KD was placed in a cubic periodic box with solvent padding on all sides (minimum 1.2 nm) using Modeller.addSolvent, neutralised, and brought to 0.15 M NaCl. To ensure consistent solvation NPT ensembles across conformations, and that conformational differences did not alter the solvent padding, the number of solvent molecules was held consistent at the maximum number of solvent molecules across all conformations. In other words, the largest possible number of solvent molecules designated by the padding cutoff was applied to all conformations. Bonds to hydrogen were constrained, water was held rigid, and hydrogen-mass repartitioning (HMR, hydrogen mass = 4.0 amu) was applied so that a 4 fs Langevin timestep could be used. Nonbonded interactions were treated with PME at a 1.0 nm real-space cutoff. Systems were energy-minimized, then assigned initial velocities from a Maxwell–Boltzmann distribution at 310 K (setVelocitiesToTemperature(310)) and propagated for 5 ns in the NPT ensemble (310 K, 1 atm) with a Monte Carlo Barostat and the Langevin Integrator with a BAOAB-like splitting (collision rate 1.0 ps^−1^, constraint tolerance 10^−5^). These equilibrated structures were launched in full on the Folding@home platform as production runs (41). Production runs were launched with 1000 NPT trajectories per starting structure (“per seed”) using OpenMM 8.0.0 with CUDA mixed precision. Integrator settings, thermostat, barostat, force field, and timestep were identical to equilibration. Example simulation scripts are provided at https://github.com/sukritsingh/distributed-sampling.

### 4.5 DFG conformational state assignment

Simulation snapshots of ABL, EGFR, and MET were assigned to DFG-in, DFG-inter, or DFG-out spatial classes by clustering the *D*_1_ and *D*_2_ distance space using HDBSCAN (83). Each snapshot was then further annotated using the Modi–Dunbrack DFG conformational nomenclature (14). In this scheme, states are labelled as BLAminus, BLAplus, ABAminus, BLBminus, BLBplus, BLBtrans, BABtrans, or BBAminus. The first three letters denote the Ramachandran basins occupied by the backbone dihedrals of the X-DFG, DFG-Asp, and DFG-Phe residues, where X-DFG is the residue immediately preceding the DFG motif. The suffix minus, plus, or trans denotes the *χ*_1_ rotamer of DFG-Phe. For each snapshot, a seven-dimensional torsion vector,

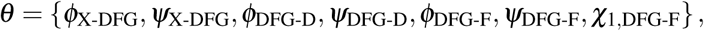

was compared with the reference centroid *θ^d^* of each Dunbrack state using the periodic cosine distance

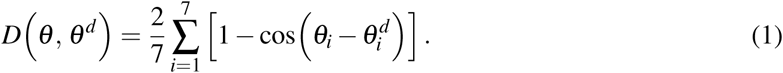

Snapshots were assigned to the Dunbrack state with the smallest distance. Snapshots for which all state-centroid distances exceeded 1 were classified as noise.

### 4.6 Markov state model construction

Trajectories shorter than 100 ns were discarded. Structural features (see Supplementary Table S2) were extracted, subjected to tICA (lag 100 ns, top 20 components), and discretized into 1000 microstates by *k*-means clustering. Transition counts were computed using effective counts with pseudo-count regularization. Reversible maximum-likelihood MSMs and Bayesian MSMs were estimated using deeptime (84). The choice of MSM parameters was validated by the implied timescale convergence (Supplementary Figs. S7, S8, S9, and S10) and kept consistent for ABL, EGFR, and MET. Macrostates were coarse-grained by PCCA+. Reactive pathways were computed using transition path theory (85). Analysis and plotting scripts are available at https://github.com/meyresearch/kinase_analysis.

### 4.7 Enhanced sampling simulations

Accelerated MD (86), Gaussian accelerated MD (87), umbrella sampling (20*×*20 grid on *D*_1_/*D*_2_), adaptive sampling (88), and metadynamics (89) were deployed on Folding@home and HPC resources targeting MET KD. Reweighting was performed by Miao cumulant expansion (aMD, GaMD) (90), MBAR (umbrella sampling) (91), or summation of Gaussians (in metadynamics) (92). All enhanced sampling methods that intake collective variables to sample/bias along used the same pair of geometric features for MET kinase, distances *D*_1_ and *D*_2_. Full details are provided in the Supplementary Methods.

## 5 Data availability

All analysis scripts and plotting scripts for MSM construction and analysis are available online at https://github.com/meyresearch/kinase_analysis. MSMs and related models are deposited on Zenodo at https://doi.org/10.5281/zenodo.20544389. The KinCore-sorted template library is hosted at https://osf.io/spu2a/. All scripts for setup and equilibration of seed structures, as well as the trimmed seed structures, are available at https://github.com/sukritsingh/kinase-seeding. Simulation setup and distributed-sampling scripts are available online at https://github.com/sukritsingh/distributed-sampling, including scripts describing simulation parameters for unbiased simulations. Scripts for the framework of adaptive sampling on Folding@home are available at https://github.com/sukritsingh/fahdaptive. The raw MD simulation trajectories for ABL, EGFR and MET comprise approximately 0.5, 1.5 and 2.3 TB of data, respectively. Owing to the size of these datasets and the lack of online storage capacity, the raw trajectories are not deposited in a public repository but are freely available upon any request.

## 6 Acknowledgements

We thank the citizen-scientists of Folding@home for donating their computing resources that enabled the distributed simulations (projects 16497–16499, 17601–17605, 17645, 17646, 17649, 17650). This work used resources from the High-Performance Computing Group at Memorial Sloan Kettering Cancer Center. The authors are grateful to the MSKCC DigITs and HPC team, especially Jamie Cheong, Lohit Valleru, and Monica Chakradeo for their assistance with high-performance computing resources. S.S. is a Damon Runyon Quantitative Biology Fellow from the Damon Runyon Cancer Research Foundation (DRQ-14-22) and acknowledges support from an NCI Pathway to Independence Award for Outstanding Early-Stage Postdoctoral Researchers (NCI K99 CA286801). The authors are thankful to John Chodera and Markus Seeliger for their helpful thoughts and feedback.

## 7 Author contributions

Y.Z., M.T.D., A.S.J.S.M., and S.S. designed the study. Y.Z. and S.S. performed simulations and analysis. S.S. developed the distributed simulation and transfer-seeding framework. M.T.D. contributed to analysis methods. M.T.D. A.S.J.S.M. and S.S. supervised the project. All authors contributed to writing the manuscript.

## 8 Competing interests

The authors declare no competing interests.

## 9 Disclaimer

The content is solely the responsibility of the authors and does not necessarily represent the official views of the National Institutes of Health.

## Notes

https://doi.org/10.5281/zenodo.20544389

https://osf.io/spu2a/

https://github.com/meyresearch/kinase_analysis

https://github.com/sukritsingh/kinase-seeding

https://github.com/sukritsingh/distributed-sampling

https://github.com/sukritsingh/fahdaptive

https://colab.research.google.com/drive/1Dcb5ZouYJQfyq1np-TBmKJE8DOAbA9dU

### Competing Interest Statement

The authors have declared no competing interest.

### Summary of Updates

1. Reframing the manuscript to focus on the *details* of activation loop dynamics based on peer review feedback 2. Fixed a typo in Figure 3D 3. Clarifying details about how enhanced sampling was conducted using the same collective variables 4. Clarifying additional details about how our models observe known states + observe new states in the same ensemble 5. Additional figures in the supplemental info. that are relevant to points 3 and 4 above 6. Additional figures in the supplemental info. that demonstrate model convergence and sufficient sampling

