## supplemental-information for "Transfer seeding reveals divergent activation ensembles across tyrosine kinases"

#### 1 Supplementary methods for enhanced sampling simulations

##### 1.1 Accelerated MD (aMD) on Folding@home

We used Hamelberg-style accelerated MD (1), in which a boost is applied to the total potential energy $V(\mathbf{x})$ whenever $V < E$:

$$V^*(\mathbf{x}) \;=\; V(\mathbf{x}) + \Delta V(\mathbf{x}), \qquad \Delta V(\mathbf{x}) \;=\; \begin{cases} \dfrac{(E - V(\mathbf{x}))^2}{\alpha + E - V(\mathbf{x})}, & V(\mathbf{x}) < E, \\ 0, & V(\mathbf{x}) \geq E, \end{cases} \tag{1}$$

so that forces are scaled by $\partial V^*/\partial V = [\alpha/(\alpha + E - V)]^2$ in the boosted region. The boost parameters $(E, \alpha)$ were chosen from a short (unbiased) calibration run: we recorded $\langle V \rangle$ over the equilibrated trajectory and set $E = \langle V \rangle + \alpha$ with $\alpha = N_{\text{atom}}/5$ for total-potential boosting. An analogous torsion-only variant was also tested (energy and $\alpha$ derived from $\langle V_{\text{tor}} \rangle$ and $3.5\, N_{\text{res}}/5$). The force modification was implemented as a `CustomIntegrator` in OpenMM that performs a standard VRORV Langevin step while replacing $\mathbf{f}$ with $\mathbf{f} \cdot [\alpha/(\alpha + E - V)]^2$ inside the boost region; the $\Delta V$ value was stored as an integrator global and dumped at every step for later reweighting (2). One thousand independent aMD replicas were run on the Folding@home from a single apo MET snapshot. Reweighting to the unbiased potential of mean force on $(D_1, D_2)$ was performed with the Miao cumulant expansion up to second order (2).

##### 1.2 Umbrella sampling on Folding@home

The $(D_1, D_2)$ plane was tiled with a $20 \times 20$ grid of independent umbrella windows, with centers $(r^0_{1,i}, r^0_{2,j})$ taken from `np.linspace(0.3, 2.0, 20, endpoint=False)` nm on each axis. In each window the system was augmented with two `CustomBondForce` harmonic biases

$$U_{\text{bias}}(\mathrm{D}_1, \mathrm{D}_2) = \tfrac{1}{2}k\,(\mathrm{D}_1 - r^0_{1,i})^2 + \tfrac{1}{2}k\,(\mathrm{D}_2 - r^0_{2,j})^2, \tag{2}$$

with force constant $k = 100\ \mathrm{kJ\,mol^{-1}nm^{-2}}$ applied between the three atom indices that define D1 and D2. Each window was equilibrated for a short NPT run and then launched as an independent Folding@home work unit using a Langevin integrator (the same VRORV Langevin kernel as above, with $\Delta V$ logging disabled). The 2D potential of mean force was obtained by MBAR reweighting over all windows (3).

##### 1.3 Metadynamics

Well-tempered metadynamics (4) was run with PLUMED 2 (5) coupled to OpenMM via the `openmmplumed` plug-in. The bias was deposited along two collective variables: $D_1$ and $D_2$, with Gaussian height 0.3 kJ $\mathrm{mol}^{-1}$ and widths $\sigma = (0.1, 0.1)$ in CV units, deposited every 10 integration steps. The simulation was run for $5 \times 10^5$ outer iterations of 10 steps each ($\sim$ 10 ns per replica) on a single HPC node. The reweighted free-energy surface was obtained by the Tiwary–Parrinello time-independent estimator (6).

##### 1.4 Gaussian accelerated MD (GaMD)

Gaussian-accelerated MD (7) uses a harmonic boost on the potential energy,

$$\Delta V(\mathbf{x}) \;=\; \begin{cases} \frac{1}{2}\,k\,(E - V(\mathbf{x}))^2, & V(\mathbf{x}) < E, \\ 0, & V(\mathbf{x}) \geq E, \end{cases} \qquad k \;=\; k_0\,\frac{1}{V_{\max} - V_{\min}}, \tag{3}$$

where $E$ is set to the running maximum of $V$ and $k_0 \in (0, 1]$ is tuned so that the standard deviation of $\Delta V$ remains below a user-specified bound (typically $\sigma_0 \leq 10\,k_{\mathrm{B}}T$). Under this boost the effective force on coordinate $q$ is $\mathbf{f}^* = \mathbf{f} \cdot [1 - k(E - V)]$, which we again implemented as a `CustomIntegrator` performing VRORV Langevin dynamics with a multiplicative force modifier and per-step logging of $\Delta V$. Boost parameters $(E, k)$ were calibrated following Miao *et al.* (7) from a short preparatory trajectory (initial collection of $V_{\min}/V_{\max}/\langle V \rangle/\sigma_V$, then adaptive update over an "equilibration" phase, then frozen for production). Independent GaMD replicas were launched on Folding@home analogously to aMD. Reweighting on $(D_1, D_2)$ used a cumulant expansion to second order (2).

##### 1.5 Distributed adaptive sampling on Folding@home

For adaptive sampling we used an adaptive seed-controller, building on previous adaptive sampling methods (8): short trajectories are launched in rounds, an initial MSM is estimated after each round, and the next round of seeds is drawn preferentially from microstates that score high on a goal function while retaining an exploration term that up-weights undersampled microstates. Specifically, after each round all currently available MET trajectories were featurized with the DFG-Phe $\zeta$-carbon distance set, projected onto a one-dimensional kinetic-map TICA space (9) (lag 10, dim 1, kinetic_map=`True`), and discretized into $k = 100$ microstates using `pyemma.coordinates.cluster_kmeans`. For each microstate we computed a goal score $g_i$ (here, $-$RMSD of the microstate centroid to a DFG-out reference) and a count score $c_i = 1/N_i$, with $N_i$ the number of frames currently assigned to microstate $i$. Both scores were min–max rescaled, and the rank was

$$s_i \;=\; (1 - w)\,\tilde{g}_i + w\,\tilde{c}_i, \tag{4}$$

with explore weight $w = 0.5$ (a companion run used $w = 1.0$, i.e. pure counts-based exploration). Microstates were sorted by $s_i$ and a cumulative-rank cutoff was used to retain the top fraction of ranks from which $n_{\text{seeds}} = 300$ structures were drawn. For each selected frame, a fresh seed was built by deserializing the FAH `system.xml.bz2/state.xml.bz2` reference files, setting the positions and box vectors to those of the chosen frame, drawing new velocities at 310 K, and writing a new `checkpointState.xml.bz2` for the next Folding@home generation. The `next_state.py` worker-side script randomly samples one of the prepared `SEEDS/*.xml.bz2` files per clone.

##### 1.6 Analysis of enhanced sampling simulations

Trajectories from all enhanced-sampling methods were imaged and aligned with `mdtraj` (10). For each frame we computed $D_1$ and $D_2$ using the same atom-index triple used by the biasing potentials and, where applicable, the DFG-Phe $\chi 1$ torsion. Two-dimensional free-energy surfaces on $D_1$ vs. $D_2$ were obtained as follows: for equilibrium seeded MD and adaptive sampling, by direct histogramming with MSM-derived stationary weights; for umbrella sampling, by MBAR reweighting across all $20 \times 20$ windows; for aMD and GaMD, by reweighting $\Delta V(t)$ with the Miao second-order cumulant expansion (2; 7); and for metadynamics, by the Tiwary–Parrinello time-independent estimator applied to the deposited Gaussians and the $(\mathrm{D}_1, \mathrm{D}_2)$ colvar trace (6). In all cases the resulting PMF $F(\mathrm{D}_1, \mathrm{D}_2)$ was expressed in $k_{\mathrm{B}}T$ units at 310 K, clipped to a common dynamic range across methods for visualization, and overlaid with the three Modi–Dunbrack reference centroids (DFG-in, DFG-inter, DFG-out). Effective sample size for the reweighted ensembles was estimated from the normalized reweighting weights $w_i$ as $N_{\mathrm{eff}} = (\sum_i w_i)^2 / \sum_i w_i^2$.

#### 2 Enhanced sampling results for MET kinase

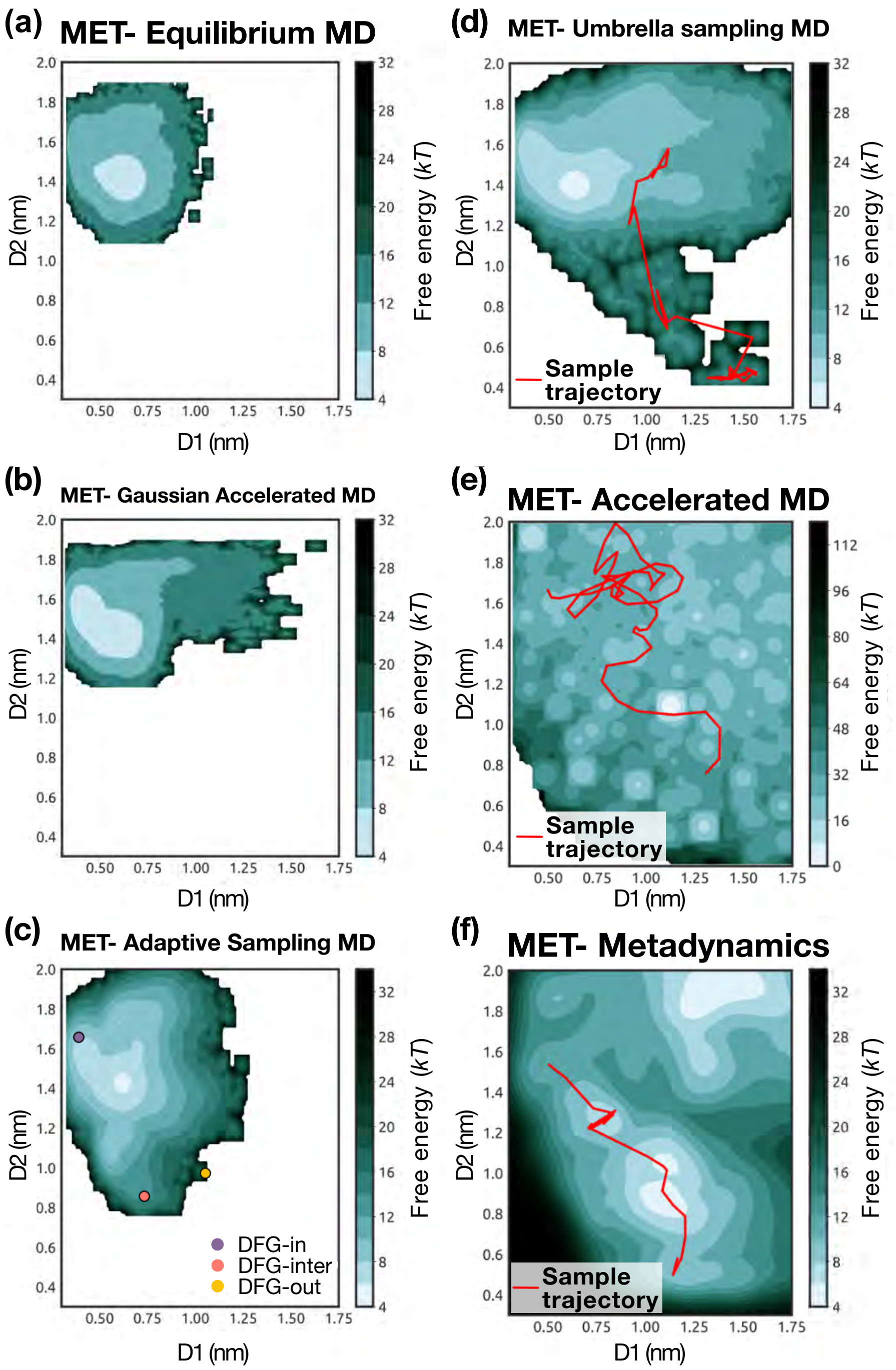


Figure S1: **Comparison of enhanced sampling methods to explore the MET DFG-transition.** Free energy surfaces projected onto $D_1$ and $D_2$ from a variety of enhanced sampling method alongside seeded equilibrium MD **(a)**. Enhanced sampling methods used and projected are distributed gaussian Accelerated MD **(b)**, distributed adaptive sampling **(c)**, distributed umbrella sampling **(d)**, distributed accelerated MD **(e)**, and metadynamics **(f)**. For adaptive sampling, representative states that discover the DFG-in (purple), DFG-inter (salmon), and DFG-out (yellow) states are marked (points). For simulations where a transition was observed in a single trajectory, a representative path through $D_1$ and $D_2$ space is drawn (red line).

For systems in which the relevant metastable states are not yet known, enhanced-sampling approaches can accelerate barrier crossing and promote exploration of rarely visited conformations. These methods alter the dynamics, energy function, or starting conditions of simulations in controlled ways to explore conformational landscapes more rapidly. Examples include metadynamics, umbrella sampling, accelerated MD (aMD), and Gaussian accelerated MD (GaMD). To avoid biasing the underlying physics that push the system away from equilibrium, adaptive sampling methods instead rapidly prioritize exploration towards an established goal by iterating between simulations from a variety of starting structures, evaluation of the exploration so far, and prioritizing new initial "promising" structures that are more likely to move towards the states of interest. Each of these methods must balance exploration with exploitation tradeoffs to rapidly explore the conformational landscape in trackable ways to recover free energy surfaces.

To quickly and efficiently identify new MET states that bridge the transition between DFG-in and DFG-out states, we deployed four enhanced sampling simulation methods in a distributed manner on the Folding@home platform: aMD, GaMD, umbrella sampling, and adaptive sampling. These approaches are history-independent, modifying only the integrator or the initial structure used for a simulation, and are thus very amenable to massively parallelized dataset generation. In conjunction, we also deployed history-dependent metadynamics simulations that, given a set of collective variables (CVs), explore conformational landscapes using history-dependent biases. This enables us to rapidly generate trajectories and assess how well these methods explore the MET kinase domain structural transition (Supplementary Fig. S1).

Of the methods tested, aMD, umbrella sampling, and metadynamics each achieved complete coverage of the DFG-in, DFG-inter, and DFG-out basins (Supplementary Fig. S1). Replicas of aMD launched from a single DFG-in structure spontaneously discovered DFG-inter and DFG-out conformations without requiring predefined collective variables. Umbrella sampling, tiling the $D_1$, $D_2$ plane with a $20 \times 20$ grid, provided dense coverage. Well-tempered metadynamics biased along $\mathrm{D}_1$, $\mathrm{D}_2$, and DFG-Phe $\chi 1$ likewise explored all three DFG basins within modest simulation time on a single HPC node. Adaptive sampling discovered the DFG-out basin through iterative rounds of short unbiased trajectories. GaMD did not achieve full coverage, with replicas remaining largely confined to the DFG-in vicinity. Although multiple methods achieved state-space coverage, the resulting free energy surfaces differ substantially across methods (Supplementary Fig. S1). Basin depths, relative free energies, and transition-region topography vary considerably, reflecting differences in bias magnitude, barrier-crossing completeness, and reweighting fidelity. A particularly illustrative case is aMD, which achieved broad state discovery but yielded a free energy surface with pronounced deep minima attributable to low effective sample size (ESS) in the reweighted ensemble. These results highlight that sampling relevant conformational states is necessary but insufficient for quantitative free energy recovery. Taken together, our results above emphasize the challenges associated with enhanced sampling simulations.

### 3 Additional supporting information

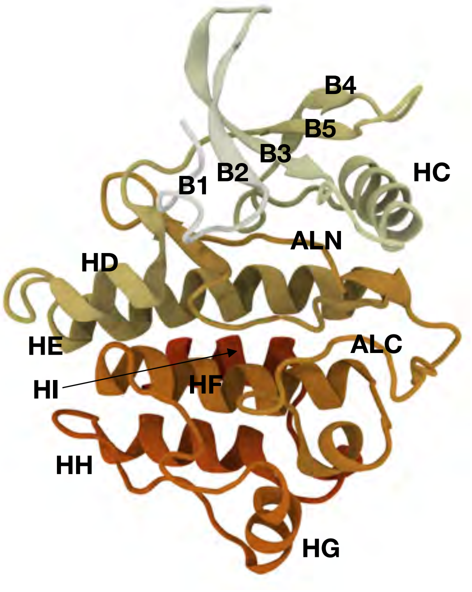


G-loop (between B1 and B2)

| | B1 | B2 | B3 | HC | B4 |
|---|---|---|---|---|---|
| ABL | 246-252 | 253-261 | 264-274 | 280-293 | 294-309 |
| EGFR | 712-722 | 723-731 | 738-748 | 756-769 | 770-784 |
| MET | 1076-1088 | 1089-1097 | 1103-1113 | 1121-1134 | 1135-1150 |

DFG (at start of ALN)

| | B5 | HD | HE | ALN | ALC |
|---|---|---|---|---|---|
| ABL | 310-320 | 321-329 | 334-357 | 374-394 | 397-413 |
| EGFR | 785-795 | 796-804 | 808-831 | 848-868 | 872-888 |
| MET | 1152-1162 | 1163-1171 | 1175-1198 | 1215-1235 | 1241-1257 |

| | HF | HG | HH | HI |
|---|---|---|---|---|
| ABL | 414-435 | 443-455 | 466-485 | 486-495 |
| EGFR | 889-910 | 918-930 | 941-960 | 961-970 |
| MET | 1258-1279 | 1287-1299 | 1310-1329 | 1330-1339 |

Figure S2: Left, representative ABL kinase domain structure annotated with segment names. Right, residue ranges corresponding to each segment in ABL, EGFR, and MET sequence alignments. Segment names follow the KinCore nomenclature, where **B** denotes $\beta$ strand, **H** denotes $\alpha$ helix, **ALN** and **ALC** denote the N- and C-terminal portions of the activation loop.

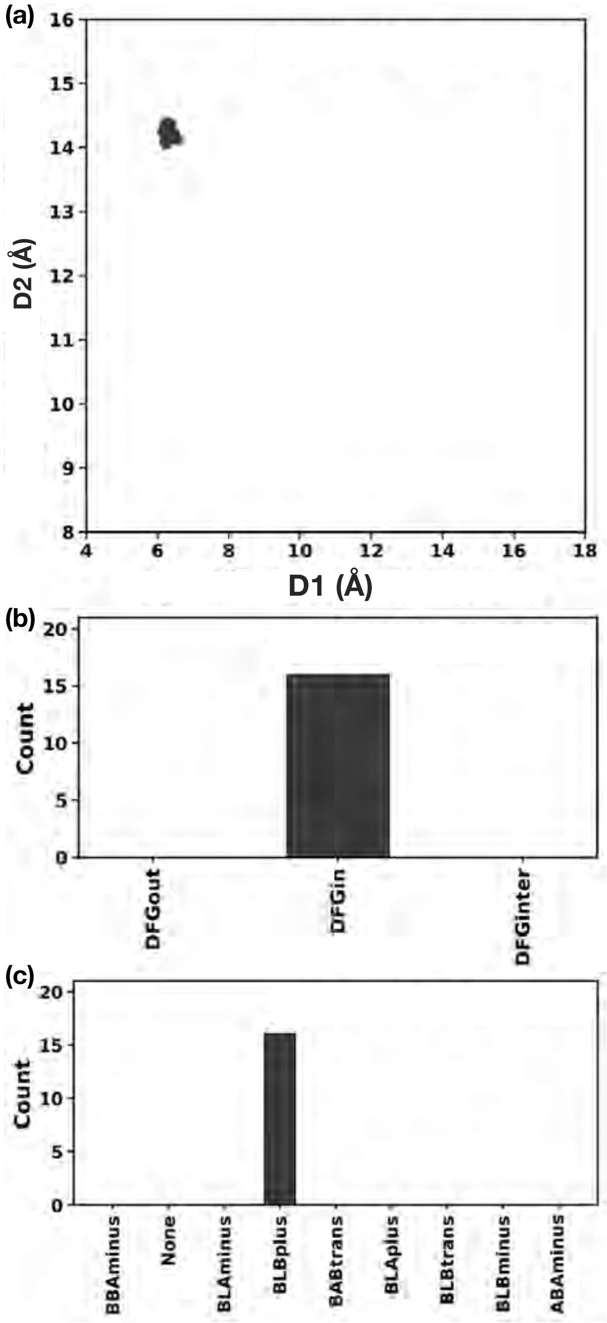


Figure S3: Template-free AlphaFold2 predictions for the MET kinase collapse to a single active-like conformation. **(a)** Two-dimensional projection of AF2 structures of MET without a template along $D_1$ ($F^{DFG}$–C$\zeta$ to $\alpha$C+4 residue M1131–C$\alpha$) and $D_2$ ($F^{DFG}$–C$\zeta$ to $\beta$3-lysine K1110–C$\alpha$). **(b)** Spatial DFG classification of AF2 generated MET structures when no template was provided. **(c)** Dihedral-based Dunbrack DFG-motif classification of each of the generated AF2 conformations without a template.

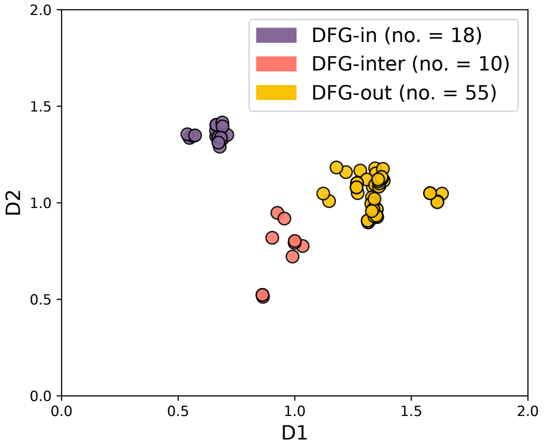


Figure S4: Crystallographic conformational landscape of ABL1 kinase domain (ABL) used as MD seeds. Distribution of all known ABL PDB structures projected onto the DFG spatial coordinates $D_1$ and $D_2$ (nm) and classified via KinCore. Each point is a PDB structure coloured by whether it is DFG-in (purple), DFG-out (yellow), or DFG-inter (salmon).

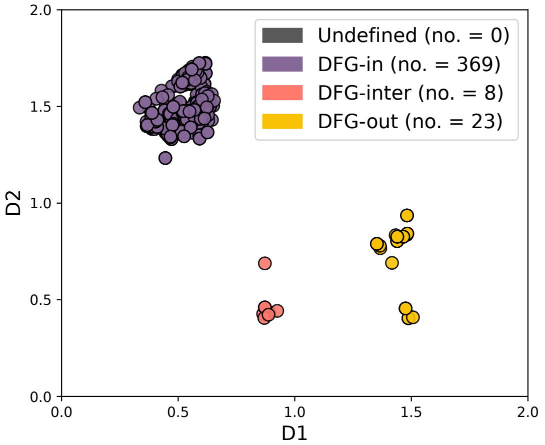


Figure S5: Crystallographic conformational landscape of EGFR kinase used as MD seeds. Distribution of all known EGFR kinase-domain PDB structures projected onto the DFG spatial coordinates $D_1$ and $D_2$ (nm) and classified via KinCore. Each point is a PDB coloured by whether the structure is DFG-in (purple), DFG-out (yellow), or DFG-inter (salmon).

Table S1: Kinase PDB IDs used for seeding MD simulations, grouped by assigned macrostate.

| Kinase | Macrostate | PDB IDs |
|---|---|---|
| ABL | S1 | 5HU9 |
| | S2 | 2HZ0 |
| | S3 | 5HU9 |
| | S4 | 1OPL, 2FO0, 2G2F, 2G2H, 2HZI, 3UE4, 4WA9, 4XEY, 4YC8, 4ZOG, 6BL8 |
| | S5 | 2F4J, 2G2I, 2GQG, 2HZ4, 2V7A, 4TWP |
| | S6 | 2E2B, 2G1T, 2HIW, 2HYY, 2HZ0, 3CS9, 3PYY, 3QRI, 3QRJ, 3QRK, 5MO4, 6NPE, 6NPU, 6NPV |
| EGFR | S1 | None |
| | S2 | 3IKA, 4G5P, 4WD5, 5UWD, 5XGN |
| | S3 | None |
| | S4 | 2JIT, 2JIU, 2RF9, 4I1Z, 4I20, 4I21, 4RIY, 4ZJV, 5HG5, 5HG7, 5HG8, 5HG9, 5U8L, 5UGA, 5UGC |
| | S5 | 1M14, 1M17, 2EB2, 2EB3, 2GS2, 2GS6, 2ITN, 2ITO, 2ITP, 2ITQ, 2ITT, 2ITU, 2ITV, 2ITW, 2ITX, 2ITY, 2ITZ, 2J5E, 2J5F, 2J6M, 2JIT, 2JIU, 2JIV, 3GOP, 3GT8, 3IKA, 3LZB, 3UG1, 3UG2, 3VJN, 3VJO, 3W2O, 3W2P, 3W2Q, 3W2R, 4G5J, 4G5P, 4I23, 4I24, 4JQ7, 4JQ8, 4JR3, 4JRV, 4LI5, 4LL0, 4LQM, 4LRM, 4R3P, 4R3R, 4R5S, 4RIW, 4RIX, 4RIY, 4RJ4, 4RJ5, 4RJ6, 4RJ7, 4RJ8, 4TKS, 4WD5, 4WKQ, 4WRG, 4ZAU, 4ZSE, 5C8K, 5C8M, 5C8N, 5CAL, 5CAN, 5CAO, 5CAP, 5CAQ, 5CAS, 5CAU, 5CAV, 5CNN, 5CZH, 5CZI, 5D41, 5EDP, 5EDQ, 5EDR, 5EM5, 5EM6, 5EM7, 5EM8, 5FED, 5FEE, 5FEQ, 5GMP, 5GTY, 5GTZ, 5HCX, 5HCY, 5HCZ, 5HIB, 5HIC, 5J9Y, 5J9Z, 5JEB, 5UGB, 5X26, 5X27, 5X28, 5X2F, 5X2K, 5XDK, 5XDL, 5XGM, 5XGN, 5Y25, 5Y9T, 5YU9, 5ZTO, 5ZWJ, 6D8E, 6JRJ, 6JRK, 6JRX, 6JWL, 6JX0, 6JX4, 6JZ0, 6LUB, 6LUD, 6P1D, 6P1L, 6S89, 6S8A, 6S9B, 6S9C, 6S9D, 6TFU, 6TFV, 6TFW, 6TFY, 6TFZ, 6TG0, 6TG1, 6V5N, 6V5P, 6V6O, 6VH4, 6VHN, 6VHP, 6WAK, 6Z4D, 7A2A, 7AEM |
| | S6 | 1XKK, 2GS7, 2RFD, 2RFE, 2RGP, 3BEL, 3GT8, 3POZ, 3W2S, 3W32, 3W33, 4HJO, 4I22, 4I24, 4ZSE, 5CNN, 5CNO, 5D41, 5GNK, 5GTY, 5X2A, 5X2C, 5X2F, 5YU9, 6DUK, 6JXT, 6P1D, 6P8Q, 6TFU, 6V5N, 6V5P, 6V66, 6V6K, 6V6O, 6WA2, 6WAK, 6Z4B, 6Z4D |
| MET | S1 | None |
| | S2 | None |
| | S3 | 3Q6W, 3R7O, 4IWD |
| | S4 | 3CTH, 3L8V, 3LQ8, 3RHK, 3VW8, 4MXC, 5HTI, 6SD9, 6SDC, 6SDD |
| | S5 | None |
| | S6 | None |
| | S7 | 5DG5 |
| | S8 | 1R0P, 1R1W, 2G15, 3CE3, 3DKC, 7B3Q, 7B3T, 7B3V, 7B3Z, 7B40, 7B41, 7B42, 7B43, 7B44 |
| | S9 | 2RFS, 2WD1, 2WGJ, 2WKM, 3A4P, 3CCN, 3CD8, 3DKF, 3DKG, 3F66, 3I5N, 3QTI, 3ZBX, 3ZC5, 3ZCL, 3ZXZ, 3ZZE, 4AOI, 4AP7, 4DEG, 4DEH, 4DEI, 4GG5, 4GG7, 4KNB, 4R1V, 4R1Y, 4XMO, 4XYF, 5EOB, 5EYC, 5EYD, 5HLW, 5HNI, 5HO6, 5HOR, 5UAB, 5UAD, 5UAF, 5YA5, 6SDE, 6UBW, 7B3W |

Table S2: Selected features of ABL, EGFR, and MET for MSM construction.

| **Feature** | **Description** | **Dimensions** |
|---|---|---|
| DFG distances | The distances of DFG-Phe-Cζ from (i) αC-Met-Cα and (ii) β3-Lys-Cα | 2 |
| DFG dihedrals | $\phi$ and $\psi$ angles of the X-DFG, DFG-Asp, and DFG-Phe, and χ1 angle of DFG-Phe converted to sin and cos values | 18 |
| αC helix distances | The distances of αC-Glu-Cδ from (i) β3-Lys-Nζ and (ii) A-loop Arg386 (ABL), Arg1227 (MET), or Lys860 (EGFR) | 2 |
| A-loop distances | Pairwise Cα distances separated by at least two residues between Asp381–Ile403 (ABL), Asp855–Asp877 (EGFR), or Asp1222–Val1247 (MET) | 210 (ABL)<br>210 (EGFR)<br>276 (MET) |

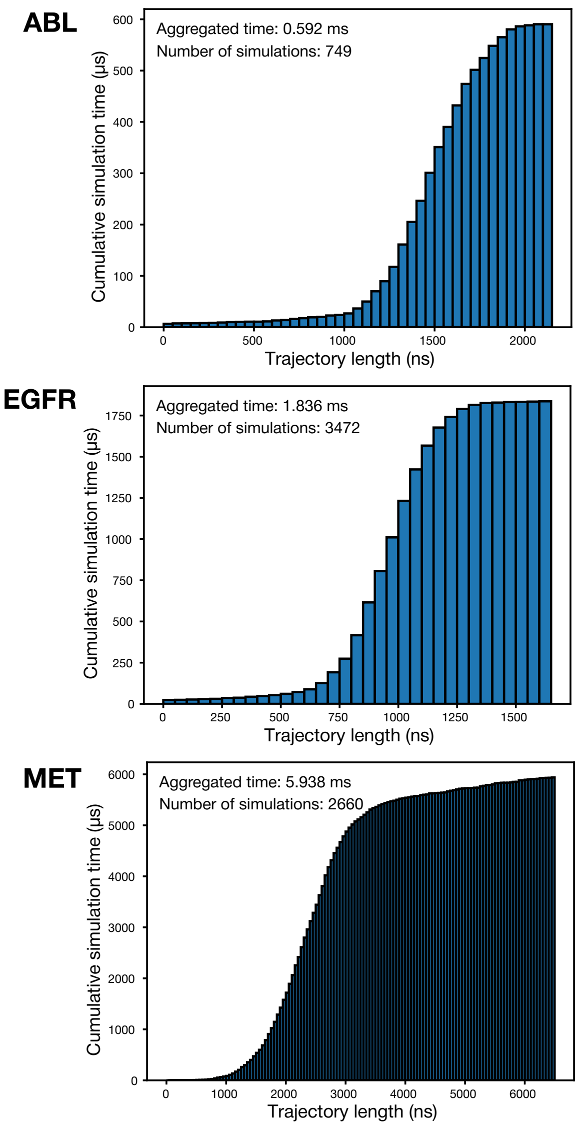


Figure S6: Cumulative simulation time for ABL, EGFR, and MET shown as a function of trajectory length binned in 50 ns.

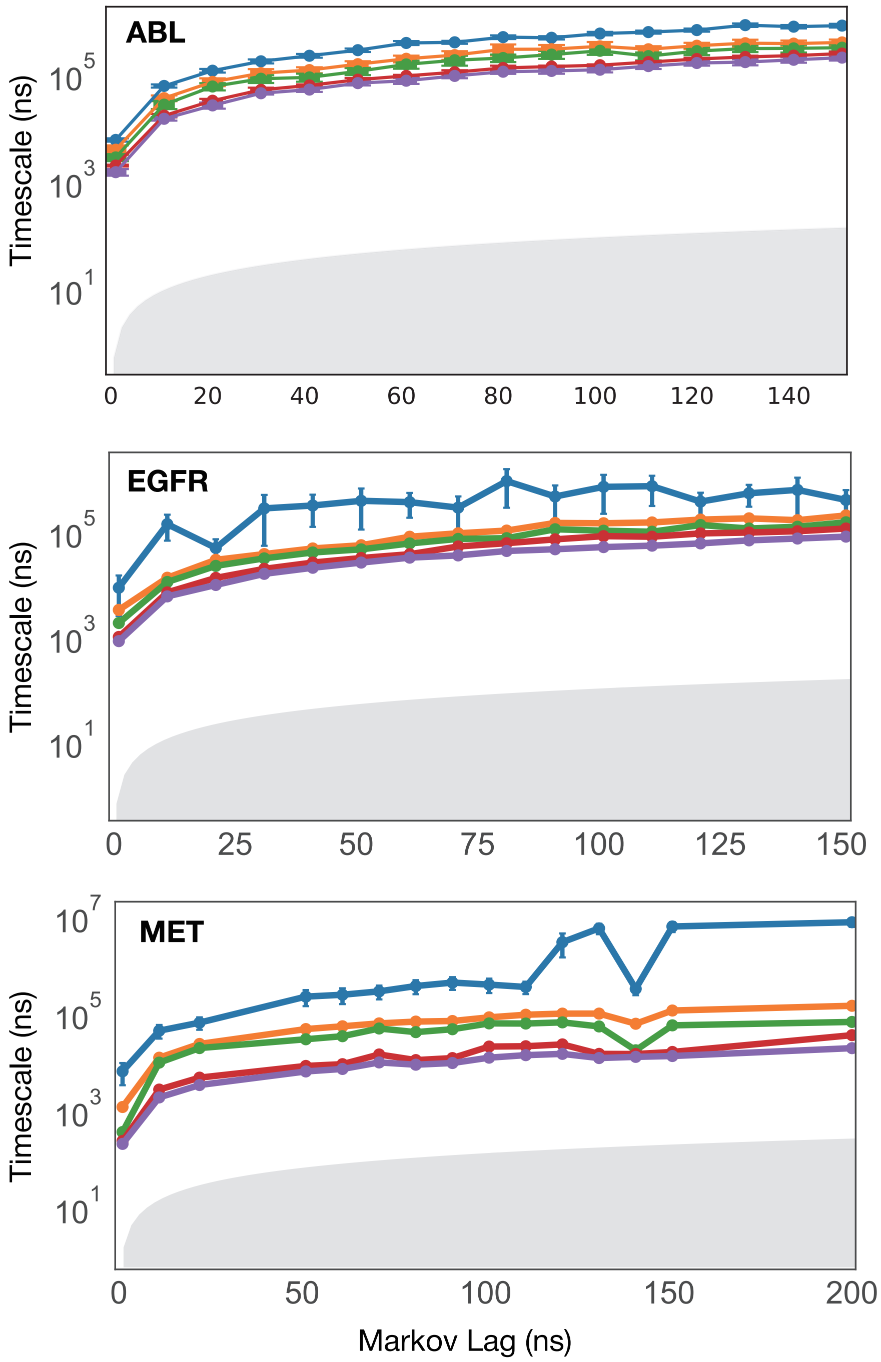


Figure S7: The five slowest timescales for ABL, EGFR, and MET estimated at Markov lag times from 1 ns to 151 ns in increments of 10 ns. The shaded area indicates the identity of timescales. The error bars show the standard deviation derived from Bayesian MSMs.

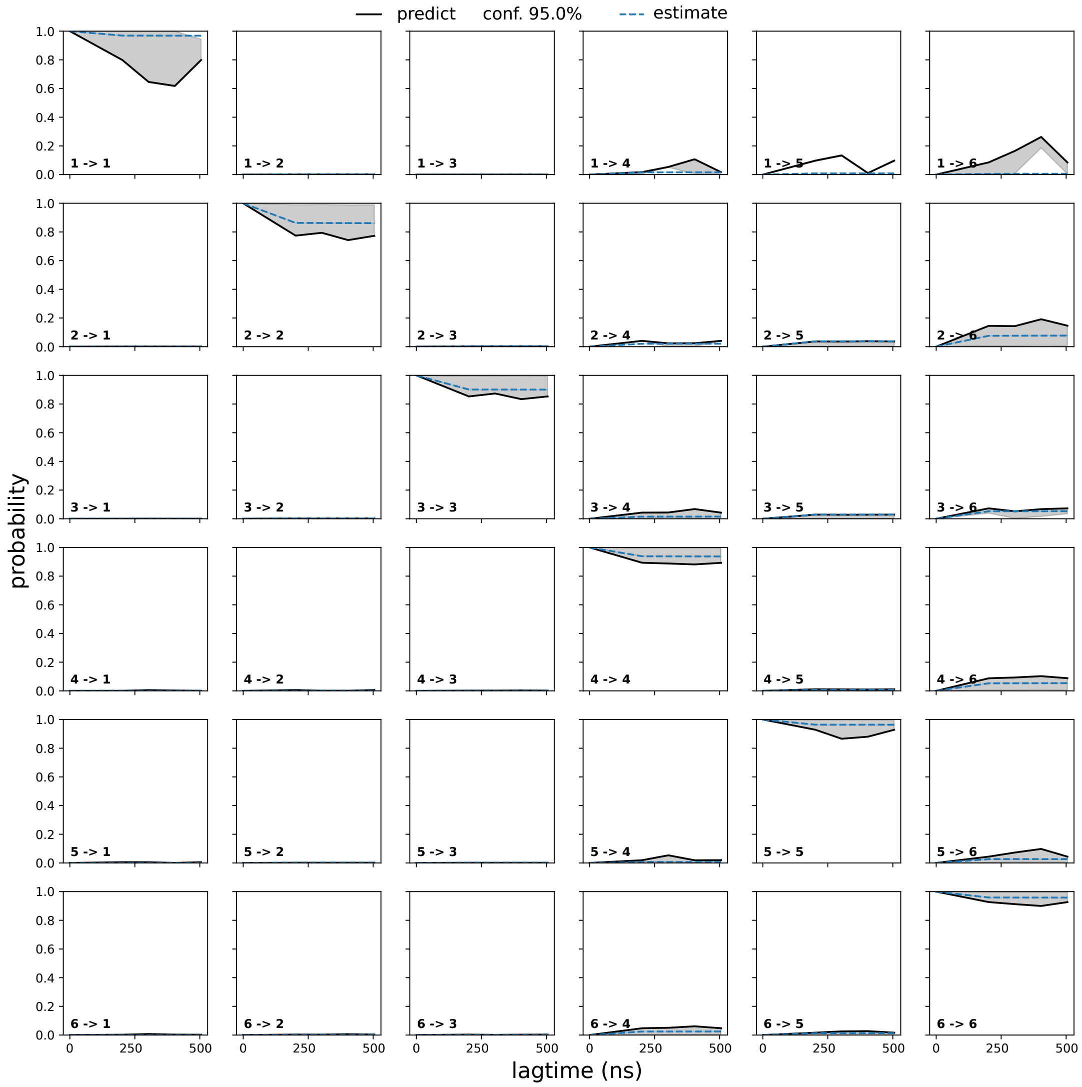


Figure S8: The Chapman-Kolmogorov tests of the ABL MSM. Transition probabilities between all pairs of macrostates are shown as a function of lag times estimated at 101, 202, 303, 404, and 505 ns. Back lines indicate the transition probabilities predicted by the MSM, the shaded regions denote 95% confident interval, and the blue dashed lines show the directly estimated transition probabilities at longer lag times. Agreement between predicted and estimated probabilities provides a test of the model's Markovian consistency.

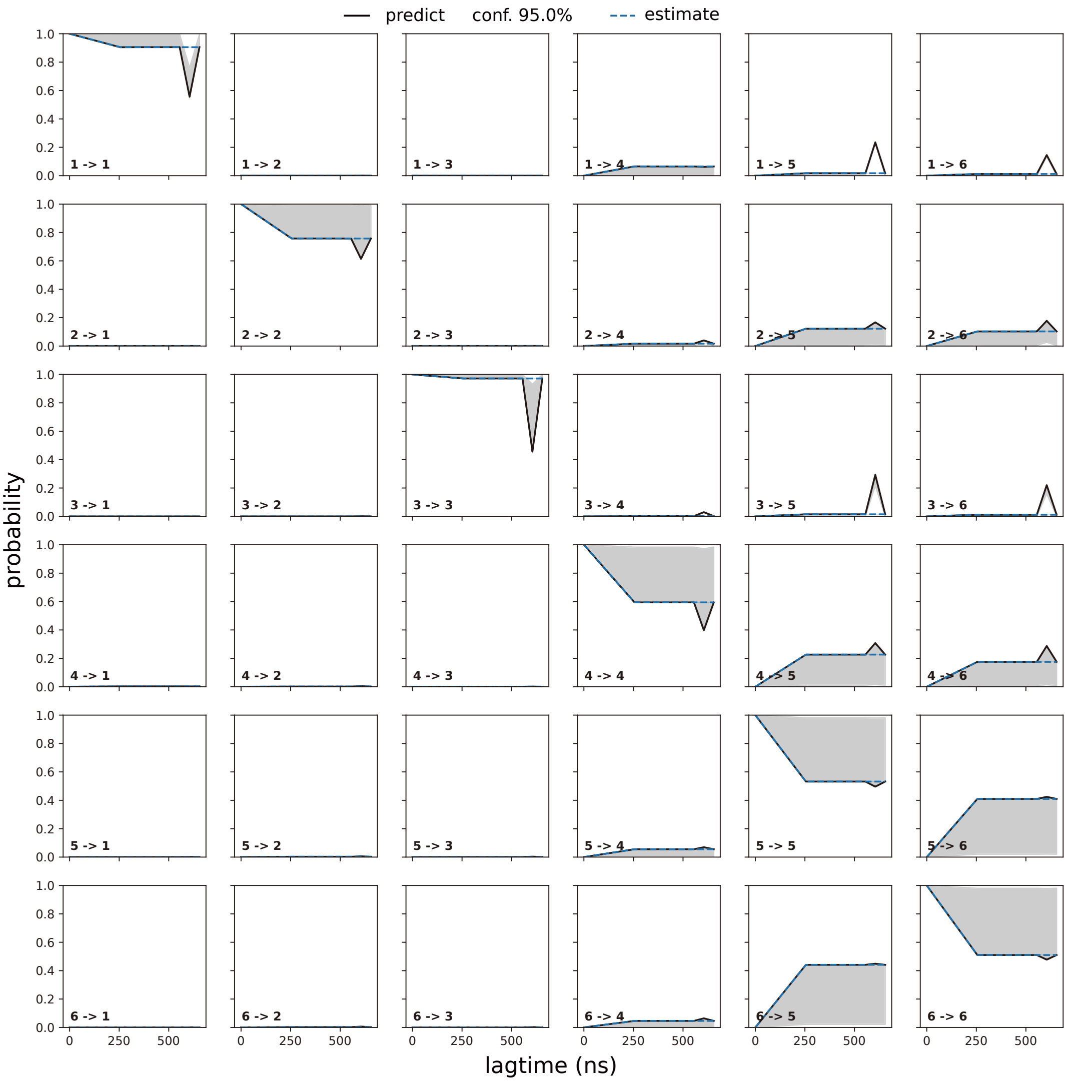


Figure S9: The Chapman-Kolmogorov tests of the EGFR MSM. Transition probabilities between all pairs of macrostates are shown as a function of lag times estimated at 101, 202, 303, 404, and 505 ns. Back lines indicate the transition probabilities predicted by the MSM, the shaded regions denote 95% confident interval, and the blue dashed lines show the directly estimated transition probabilities at longer lag times. Agreement between predicted and estimated probabilities provides a test of the model's Markovian consistency.

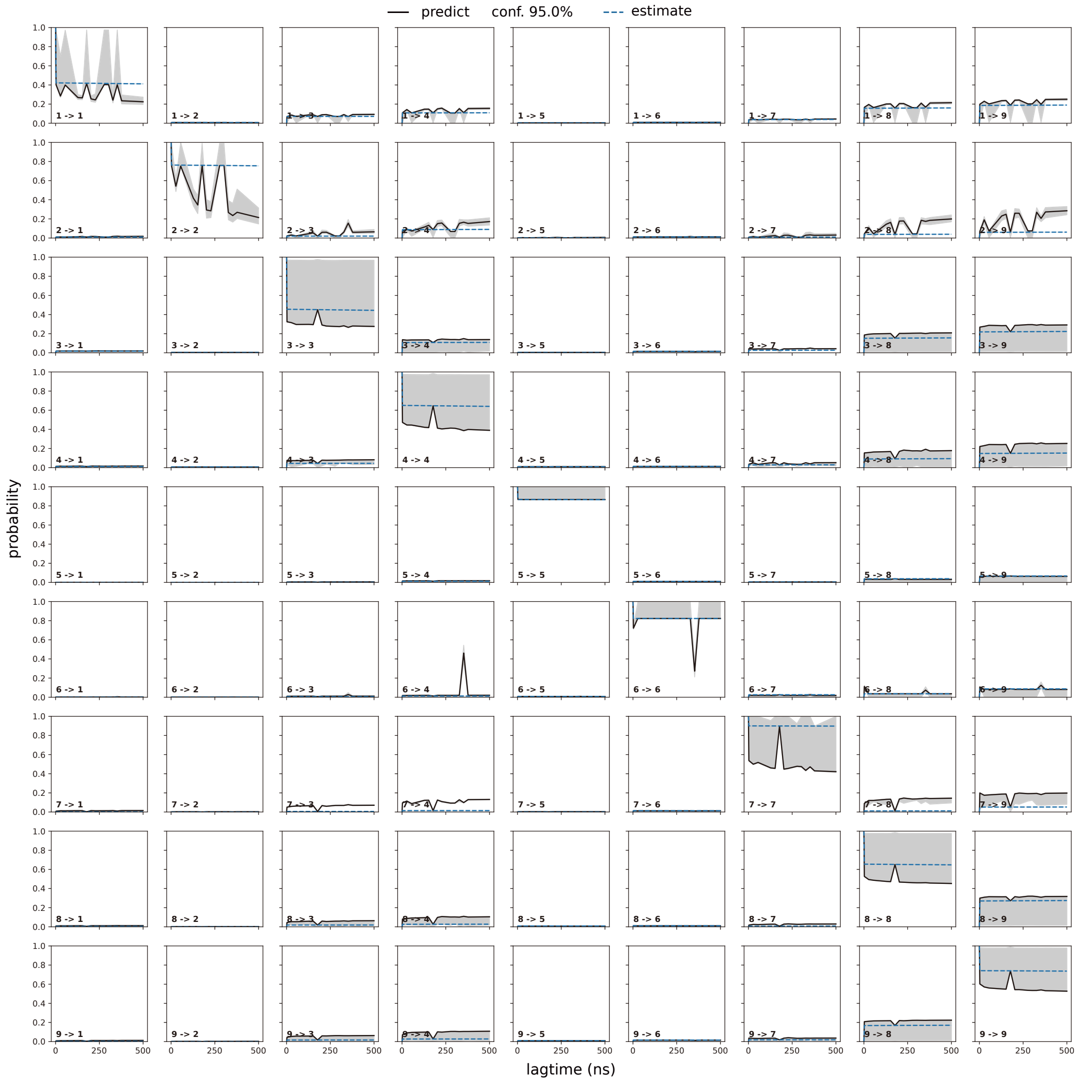


Figure S10: The Chapman-Kolmogorov tests of the MET MSM. Transition probabilities between all pairs of macrostates are shown as a function of lag times estimated at 101, 202, 303, 404, and 505 ns. Back lines indicate the transition probabilities predicted by the MSM, the shaded regions denote 95% confident interval, and the blue dashed lines show the directly estimated transition probabilities at longer lag times. Agreement between predicted and estimated probabilities provides a test of the model's Markovian consistency.

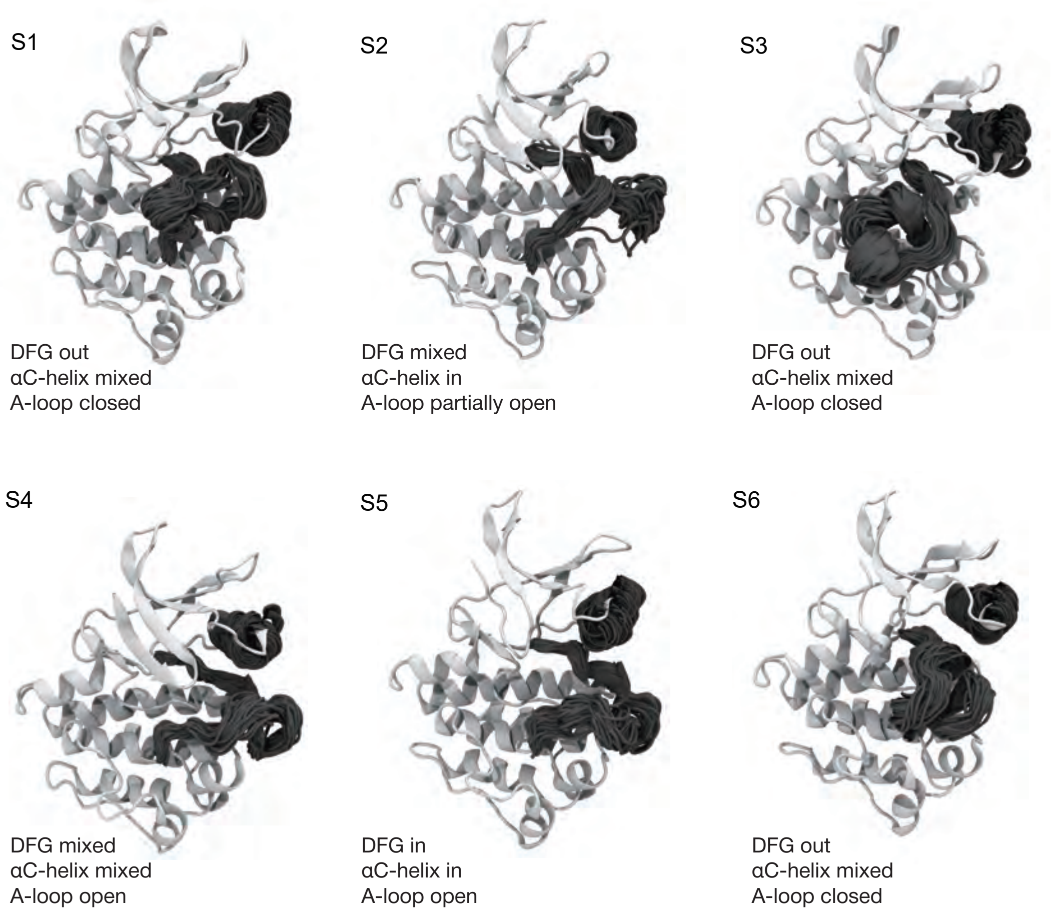


Figure S11: A-loop and $\alpha$C helix (shaded regions) structural ensembles for ABL macrostates. The corresponding macrostate is indicated (top left above each structure).

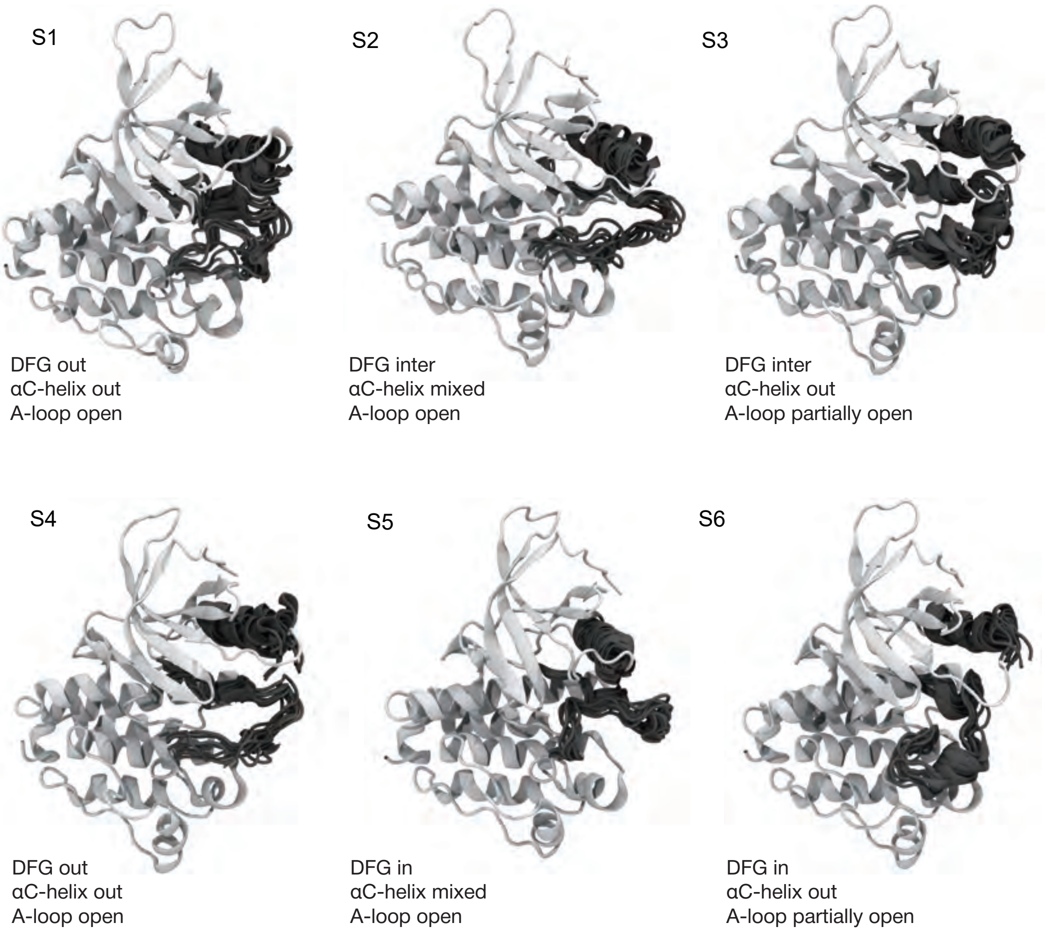


Figure S12: A-loop and $\alpha$C helix structural ensembles for EGFR macrostates. The corresponding macrostate is indicated (top left above each structure).

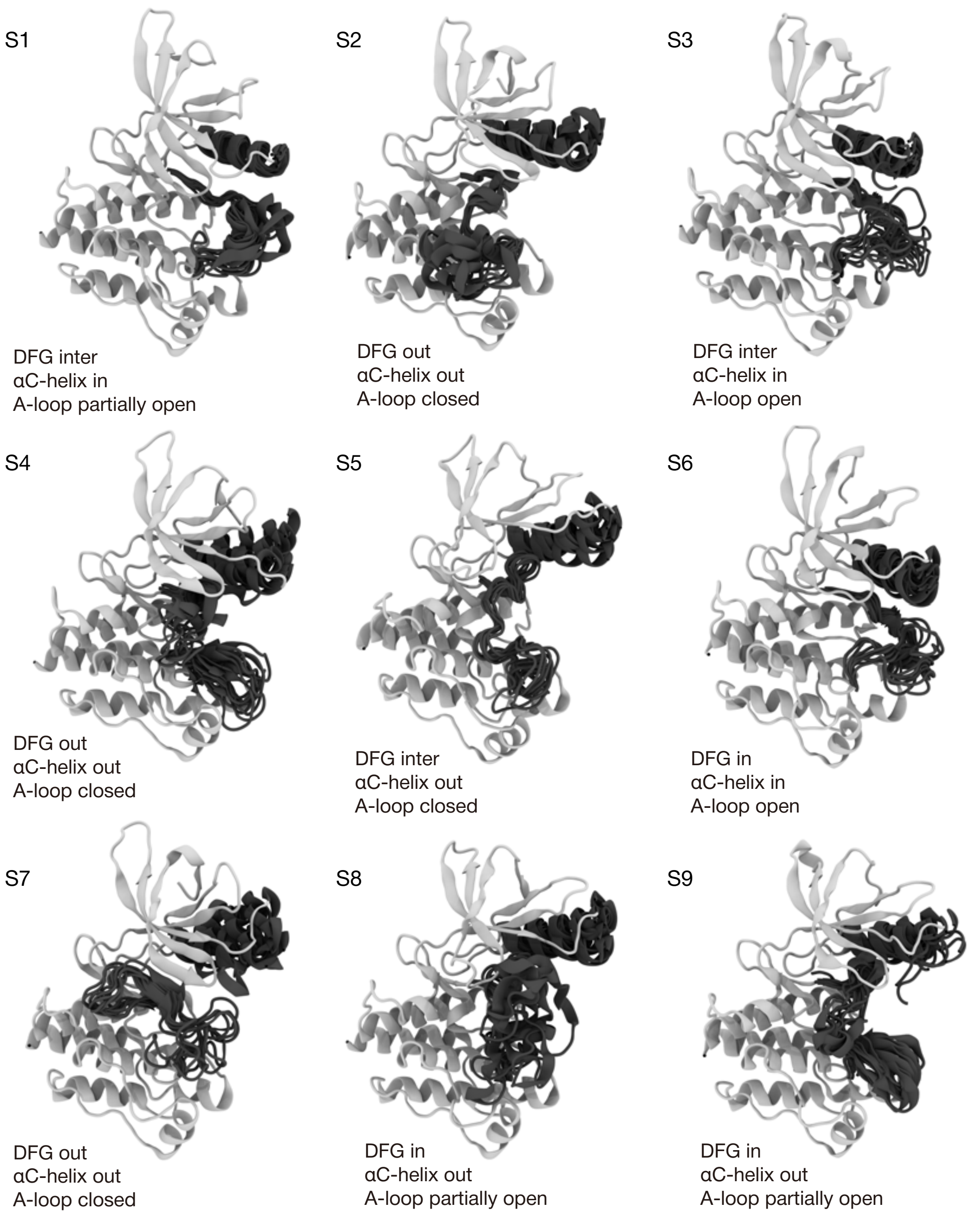


Figure S13: A-loop and $\alpha$C helix structural ensembles for MET macrostates. The corresponding macrostate is indicated (top left above each structure)

(a)

(b)

(c)

(d)

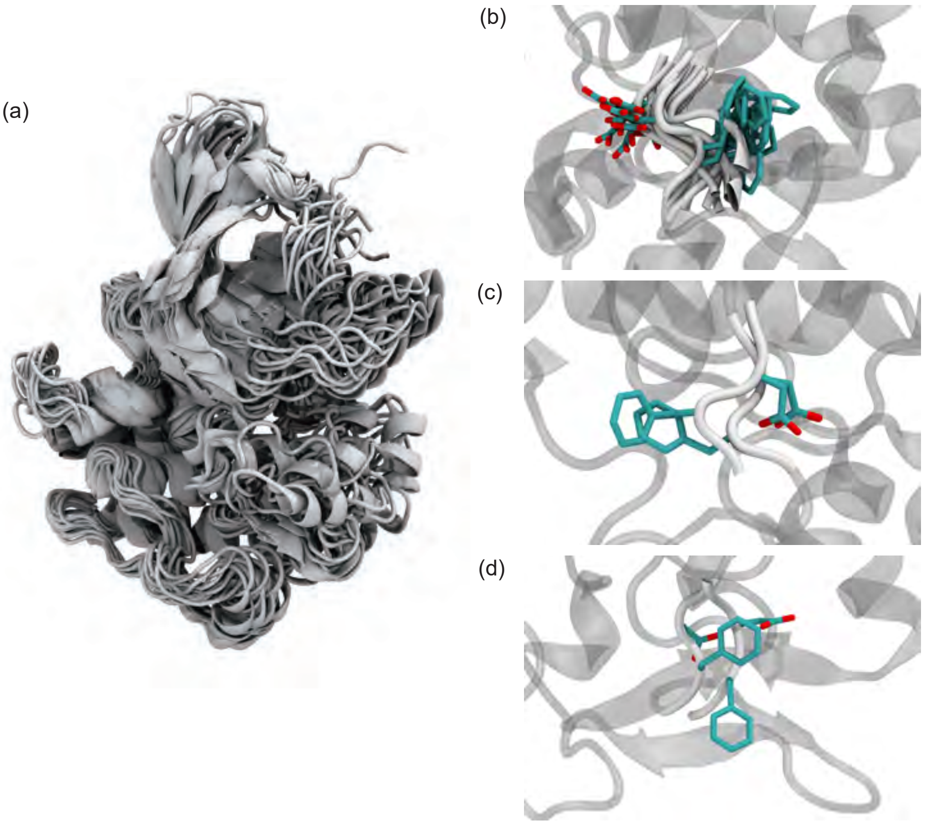

Figure S14: AF2 seeded MET conformations in different Dunbrack states derived using transfer seeding. (a) Structural overlay of all 16 seeded conformations. (b, c, d) The DFG motifs of structures seeded in DFG-in (b), DFG-inter (c), and DFG-out (d).

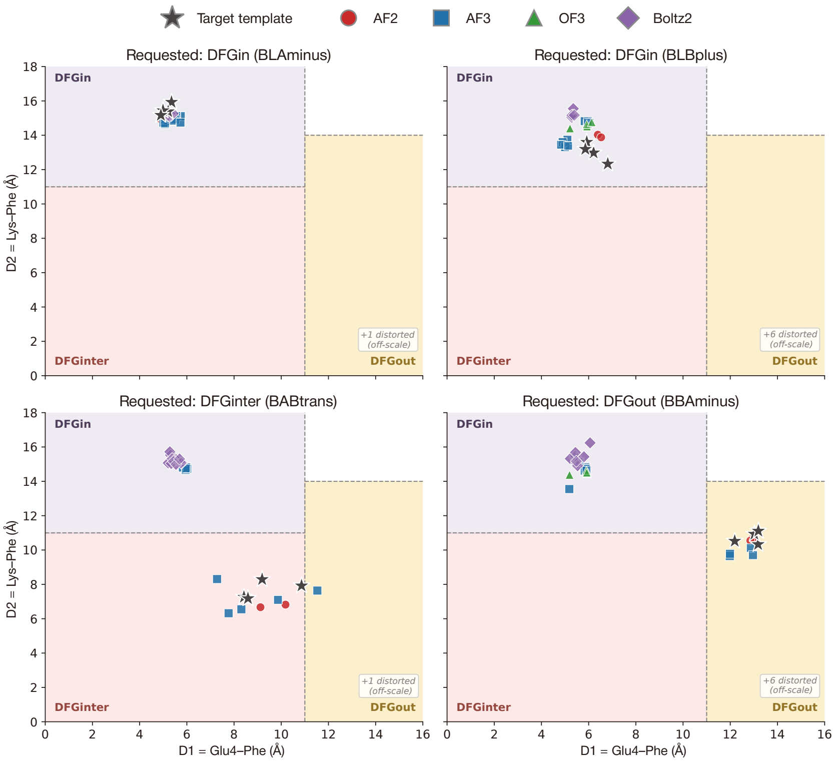


Figure S15: MET structures predicted using transfer-seeding with a variety of co-folding models given a set of templates (gray star). All structures are projected onto the same $D_1$ vs. $D_2$ access for comparison for one of four given templates: BLAminus (top left), BLBplus (top right), BABtrans (bottom left), and BBAminus (bottom right). Co-folding models tested were AlphaFold2 (red circle), AlphaFold3 (blue squre), OpenFold3 (green triangle), and Boltz-2 (purple diamond), each of which were provided with the same structural templates (gray star).

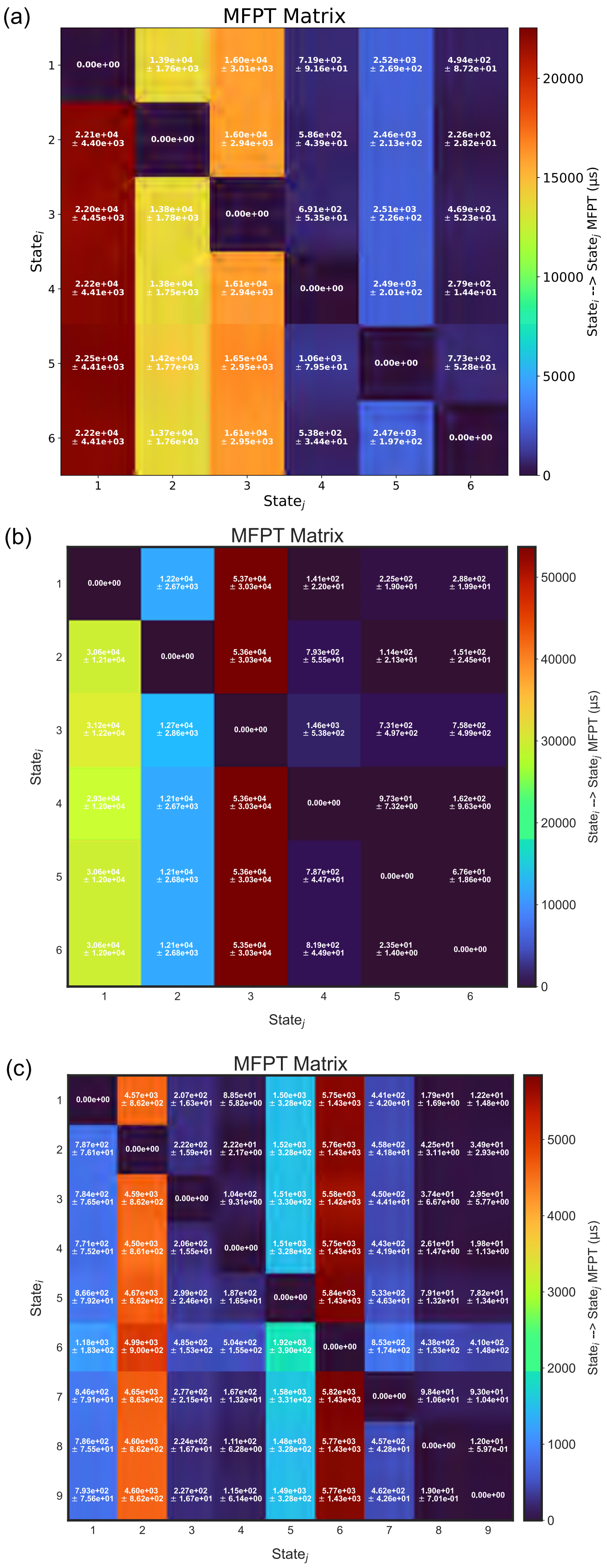


Figure S16: The mean first passage times between metastable macrostates for (a) ABL, (b) EGFR, and (c) MET. The color of the heatmap is indicative of the MFPT (colorbar, right).

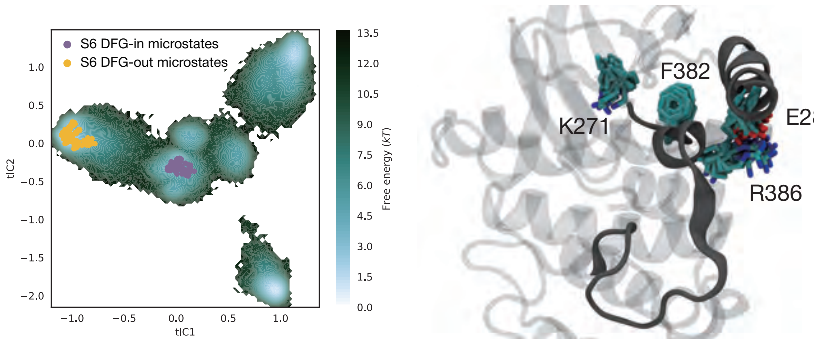


Figure S17: Projecting microstates that represent ABL microstate 6.The DFG-in BLBplus and DFG-out BBAminus microstates of ABL macrostate 6 (left). A zoomed-in view of the A-loop-Arg-αC-Glu salt bridge of the DFG-in BLBplus (SRC-like inactive) substate (right).

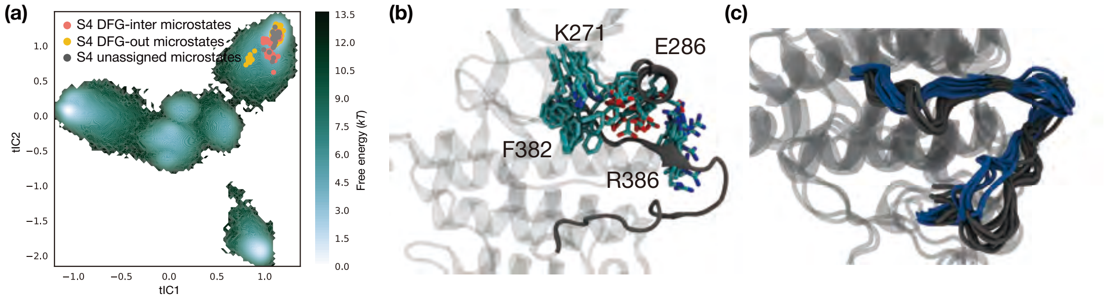


Figure S18: The microstates that make up ABL macrostate 4(a) The DFG-inter BABtrans, DFG-out BBAminus, and unassigned microstates of ABL macrostate 4. (b) A zoomed-in view of the intermittently formed $\beta$3-Lys αC-Glu salt bridge of ABL macrostate 4. (c) Comparison of ABL A-loop conformations between macrostate 4 (blue) and macrostate 5 (grey).

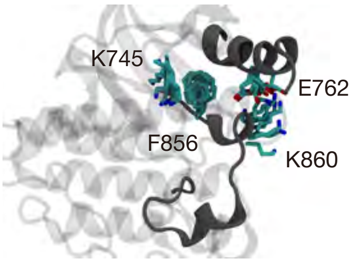


Figure S19: A zoomed-in view of the A-loop-Lys αC-Glu salt bridge of the EGFR SRC-like inactive macrostate 6.

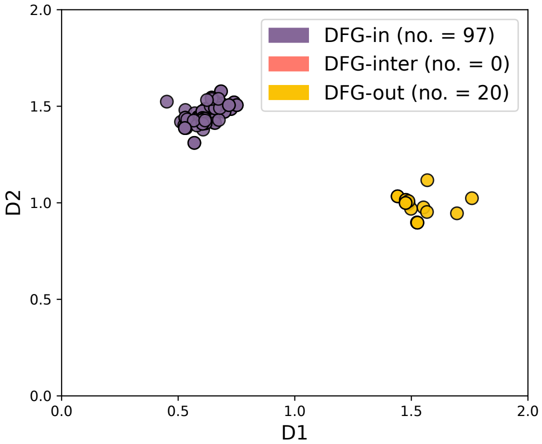


Figure S20: Crystallographic conformational landscape of MET kinase used as MD seeds. Distribution of all known MET kinase-domain PDB structures projected onto the DFG spatial coordinates $D_1$ and $D_2$ (nm) as classified via KinCore. Each point is a PDB coloured by whether the structure is DFG-in (purple), DFG-out (yellow), or DFG-inter (salmon).

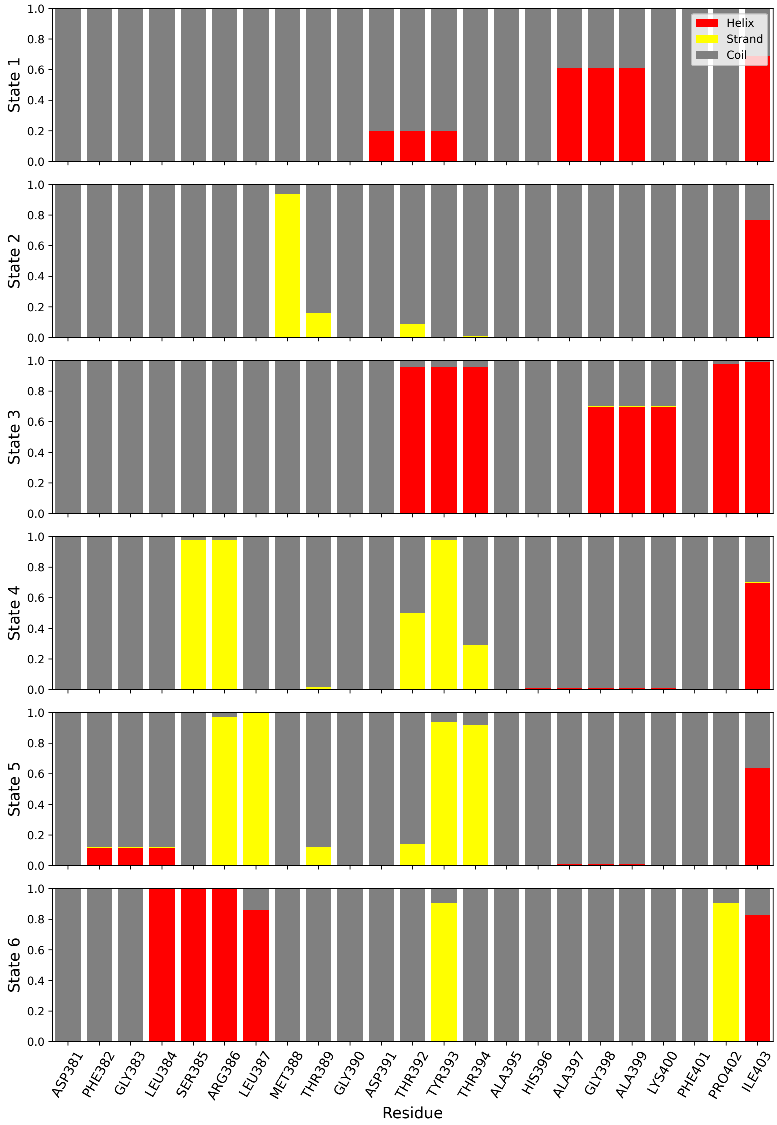


Figure S21: Secondary structure propensities of ABL A-loop subset (Asp381–Ile403) across each of the six macrostates, showing the fractional occupancy of DSSP-assigned elements for each residue. For each state (rows), each residue (column) is classified for the proportion of time spent as a helix (red), a coil (gray) or a $\beta$-strand (yellow).

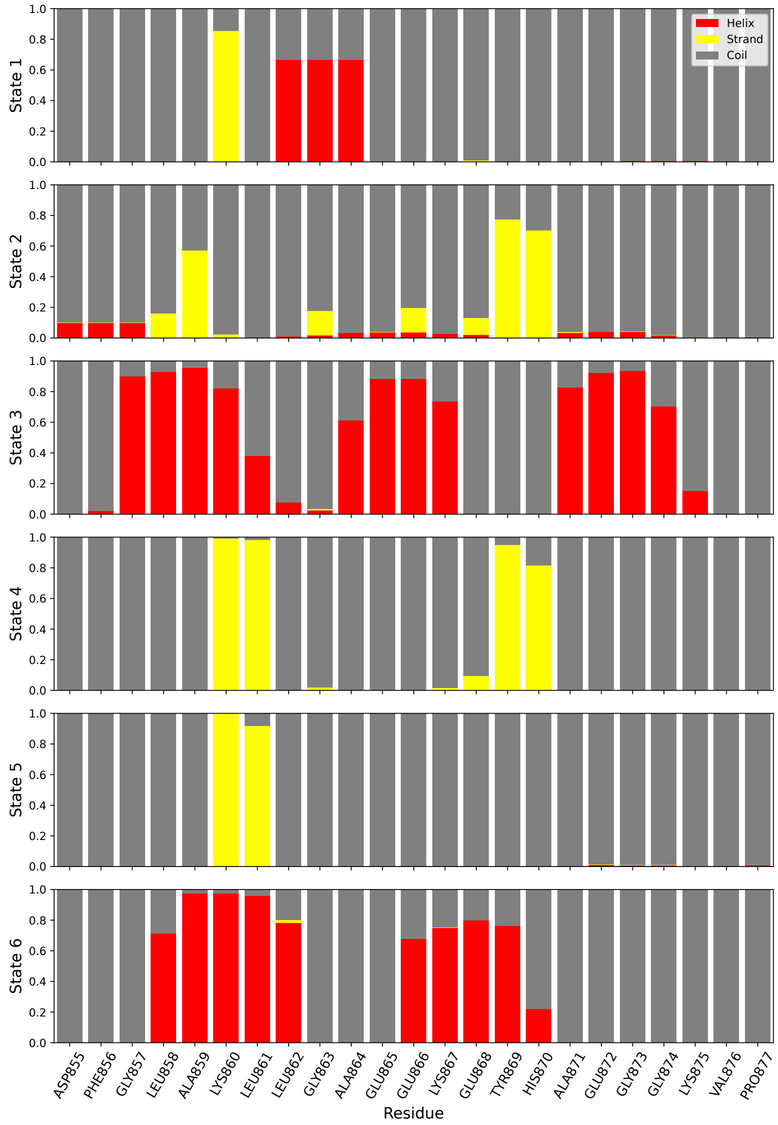


Figure S22: Secondary structure propensities of EGFR A-loop subset (ASP855–PRO877) across each macrostates, showing the fractional occupancy of DSSP-assigned elements for each residue. For each state (rows), each residue (column) is classified for the proportion of time spent as a helix (red), a coil (gray) or a $\beta$-strand (yellow).

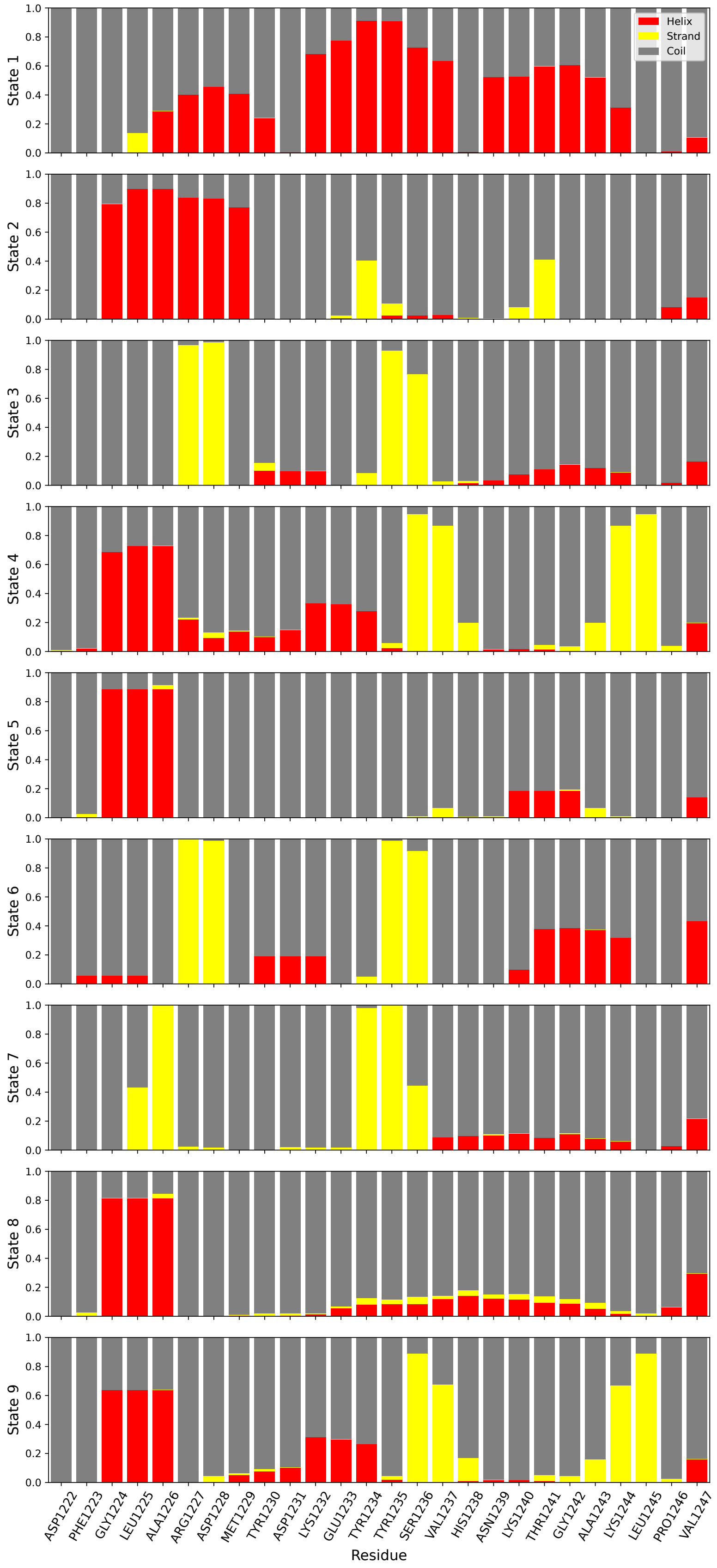


Figure S23: Secondary structure propensities of MET A-loop subset (Asp1222–Val1247) across each macrostate, showing the fractional occupancy of DSSP-assigned elements for each residue. For each state (rows), each residue (column) is classified for the proportion of time spent as a helix (red), a coil (gray) or a $\beta$-strand (yellow).

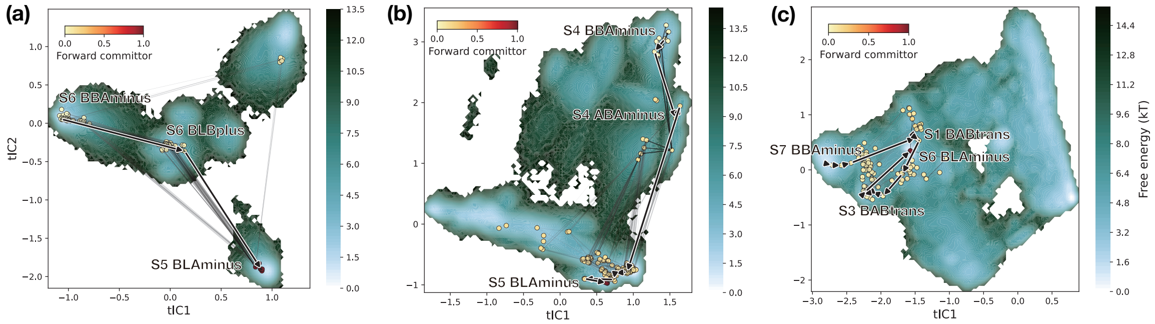


Figure S24: The top 100 highest flux pathways of (a) ABL, (b) EGFR, and (c) MET transitioning from a dominant BBAminus to BLAminus ensemble. Each line's thickness is proportional to its net transition flux. The highest flux pathway is highlighted (outlined arrows). The relevant microstates are shown and coloured by their forward committor probabilities (colorbar, top left).

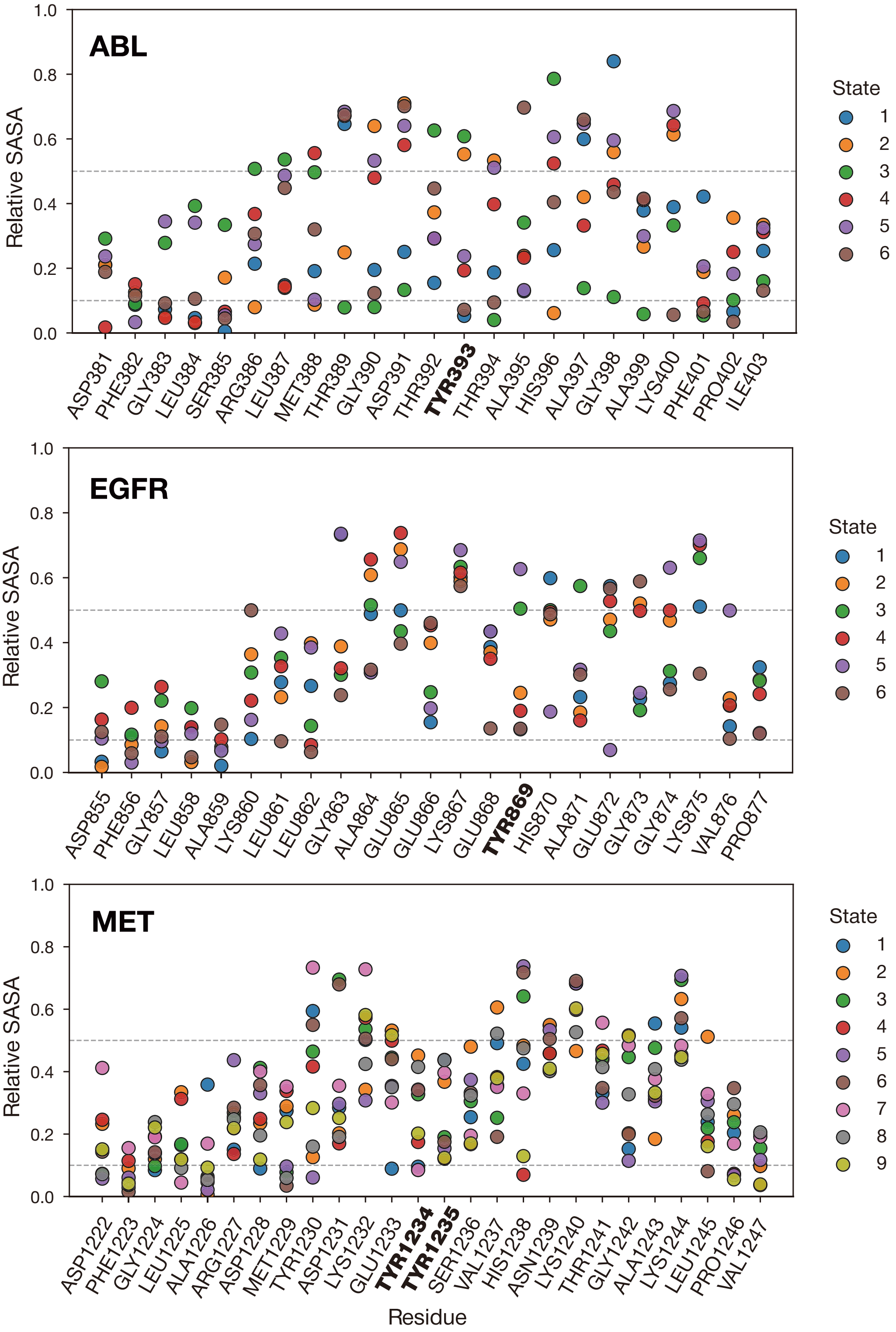


Figure S25: Residue-wise relative solvent-accessible surface area (rSASA) of the activation loops for ABL (top), EGFR (middle), and MET (bottom). Absolute SASA values were normalized by the residue-specific maximum accessible surface areas reported by Tien et al. (11), and per-residue rSASA values are reported per state (colors, right). Grey dashed lines correspond to relative SASA values of 0.1 and 0.5 to indicate complete burial or complete solvent exposure. Tyrosine phosphorylation sites reported over 50 times in PhosphoSitePlus are highlighted in bold (12).
